# Polymer of methyl malonic acid suppress inflammation by downregulating IL-2 through ROS overproduction

**DOI:** 10.64898/2026.01.11.698915

**Authors:** Sanmoy Pathak, Madhan Mohan Chandra Sekhar Jaggarapu, Taravat Khodaei, Abhirami Thumsi, Joel P. Joseph, Abhinav P. Acharya

## Abstract

Metabolites belonging to the propionate metabolism pathway can regulate immune cell responses in the context of autoimmunity and chronic inflammation. Methyl malonic acid (MMA), a metabolite in this pathway is known to cause dysregulation of T cell oxidative phosphorylation (OXPHOS) and downregulating pro inflammatory T cell effector functions. However, the effects of MMA on T cell signaling and T cell activation is not clearly known. Furthermore, since MMA is a small molecule, using it in the context of therapy remains a problem. It gets metabolized in a short time and millimolar concentrations are required to get effective results. This work describes a novel polymer, 1,6 MMA, synthesized using 1,6 Hexane-diol and MMA, which helps in slow, steady and continuous release of the small molecule. Doses in micromolar ranges generate long lasting and robust immunosuppression of activated T cells via an IL2 dependent mechanism in both human and mice T cells without causing non-specific toxicity. This causes a dysregulated expression of pSTAT5 which eventually enhances BLIMP1 mediated T cell apoptosis. Finally, 1,6 MMA mediated T cell suppression is caused due to increase in mitochondrial ROS production. Extrapolation of our findings in-vivo showed the polymer inhibited autoreactive T cell responses in mice with collagen induced arthritis (CIA). Overall, 1,6 MMA, a novel metabolite polymer, has major therapeutic potential in combating inflammatory disorders.

## 1. Introduction

Rheumatoid Arthritis (RA) is considered as a systemic autoimmune disease which is characterized by the development of inflammatory arthritis causing swelling of the joints which causes long lasting deformity^1^. The chronic inflammation associated with this disease could be triggered due to genetic and environmental factors^2,3^. Understanding the immunological mechanisms that enhance symptoms of this disease is complex. Complete dysregulation of both the innate and adaptive immune cell responses are observed at the site of inflammation^4^. Innate immune cells such as macrophages produce various proinflammatory cytokines such as TNF, IL6, IL1, IL15 that promote tissue inflammation^5–8^. On the other hand, adaptive immune cells such as T cells and B cells respond aggressively to antigenic stimuli such as citrullinated proteins^9,10^. Dendritic cells process these modified proteins, express using MHC which is recognized by T cells via the TCR-MHC interaction^11^. This leads to the development of type1 helper T cells (Th1) and type 17 helper T cells (Th17) which are one of the primary reasons for the development of chronic inflammation^12,13^. Current therapies that are used for the treatment of RA such as methotrexate, TNF inhibitors, co-stimulation blockers (abatacept) are often just temporary fixes with serious side effects such as opportunistic infection and hepatotoxicity^14–16^.

Therefore, alternative approaches to treat such diseases is of great importance. One such approach is targeting immunometabolism. Both proinflammatory and anti-inflammatory responses generated by immune cells have certain energy requirements. Metabolic pathways such as glycolysis, oxidative phosphorylation, and fatty acid oxidation provides the fuel to generate these responses for specific functions^17^. For example, proinflammatory T cell responses are largely dependent on glycolysis for their energy needs. In such a scenario, T cells undergo a phenomenon called Warburg effect where there is a dramatic shift in metabolic requirements from oxidative phosphorylation to glycolysis^18^. Alternatively, regulatory T cells (Tregs) that suppress excess proinflammatory responses rely on fatty acid oxidation and oxidative phosphorylation for their suppressive functions^19^. Therefore, tweaking metabolic signatures to maintain immune homeostasis is a great alternative approach to treat RA.

Metabolites, of mentioned pathways, modified in various forms, have previously been used to treat both autoimmune diseases and cancer. Modified metabolites in the form of polymers improved drug delivery and stability of the metabolite to enhance immunomodulation. Recent work from the lab showed the immunomodulatory effects of polyethylene succinate nanoparticles (PES NPs) in enhancing proinflammatory M1 macrophage responses thereby reducing melanoma tumor growth in mice. This study showed improved immunotherapy against melanoma when PES NPs were combined with anti-PD1^20^. Another study using polymeric nanoparticles of alpha ketoglutarate (paKG NPs) demonstrated its efficacy in reducing Th17 and enhancing Treg responses in mouse model of arthritis^21^. Methylmalonic acid (MMA) is a metabolite that is generated during the catabolism of odd-chain fatty acids and branched chain amino acids^22^. This metabolite is therefore a part propionate metabolism pathway where the end-product, succinyl-CoA enters the tricarboxylic acid (TCA) cycle^22^. Enhanced accumulation of MMA causes mitochondrial dysfunction and anergy of CD8+ T cells^23^. This metabolite specifically accumulates in tumor microenvironments causing dampening of anti-tumor T cell responses^23^. Therefore, MMA has the potential to be used as an immunosuppressive metabolite in the context of uncontrolled inflammatory responses. In this study, our primary goal was to understand whether MMA had the potential to be developed as a novel immunotherapy to reduce symptoms of RA. We specifically engineered novel polymers of MMA with 1,6 hexanediol (1,6 MMA) using a poly condensation reaction. Initial confirmation and characterization of the structure was done using FTIR and NMR studies. We initially tested its effect invitro on both CD4+ and CD8+ T cell activation where we focus our findings on the IL2 signaling pathway and CD25 expression. Next, molecular targets such as pSTAT5 and BLIMP-1 expressions were identified to understand the changes in gene regulation and downstream cell signaling. Finally, metabolic signatures were studied to understand the influence of MMA on oxidative phosphorylation and the subsequent generation of reactive oxygen species (ROS) which has the capability of causing dysregulation of proinflammatory immune cell responses. We extrapolated our findings in CIA mouse model of arthritis. The idea was to understand the in-vivo therapeutic effect of the polymer on disease symptoms and whether it was implemented by a T cell dependent mechanism. Finally, to deliver a clinical statement where we performed studies of 1,6MMA on human CD3+ T cell activation and conducted proteomic analysis to look at common molecular mechanisms. This holistic study would therefore help us understand the mechanisms by which 1,6 MMA polymer could reduce symptoms of RA and be used as a novel therapeutic strategy.

## 2. Results

### 2.1 Synthesis and characterization of novel 1,6 MMA polymer in vitro and invivo

Sustained levels of MMA are required to elicit the intended immunomodulatory response. Although esterification enhances MMA membrane permeability, achieving effective in vivo concentrations would necessitate frequent administration of large doses due to rapid diffusion and clearance associated with its low molecular weight. To address this limitation, we synthesized degradable MMA-based polymers designed (**Fig 1A**) to provide prolonged release following a single administration. In particular, MMA was polymerized with 1,12-dodecanediol, or 1,10 decanediol, or 1,8 octanediol or 1,6 hexanediol to yield polyesters with a number-average molecular weight (*Mn*) ranging from of 3.8–5.1 kDa (**Table S1**), as characterized by gel permeation chromatography (GPC). Moreover, the generation of esters was confirmed via Attenuated Total Reflectance Fourier Transform Infrared Spectroscopy (ATR-FTIR, 1735-1750 cm^-1^) (**Fig 1B**). Also, the presence of both MMA and the aliphatic carbon chain was determined using 1H NMR (characteristic downfield shift of the diol methylene protons **δ** = 4.15 ppm and the methine groups of the methylmalonic acid **δ** = 3.45 ppm) (**Fig 1C**). Subsequently, wanted to understand the effective concentration of MMA release in-vivo in mice. **Fig S1** demonstrated the stability of 1,6 MMA as we observed detectable amounts of MMA released in the blood till 21 days of a single injection of 50μg polymer via the intraperitoneal (i.p.) route. We observed the highest concentration of MMA release at 24 hours post injection.

**Figure 1:**
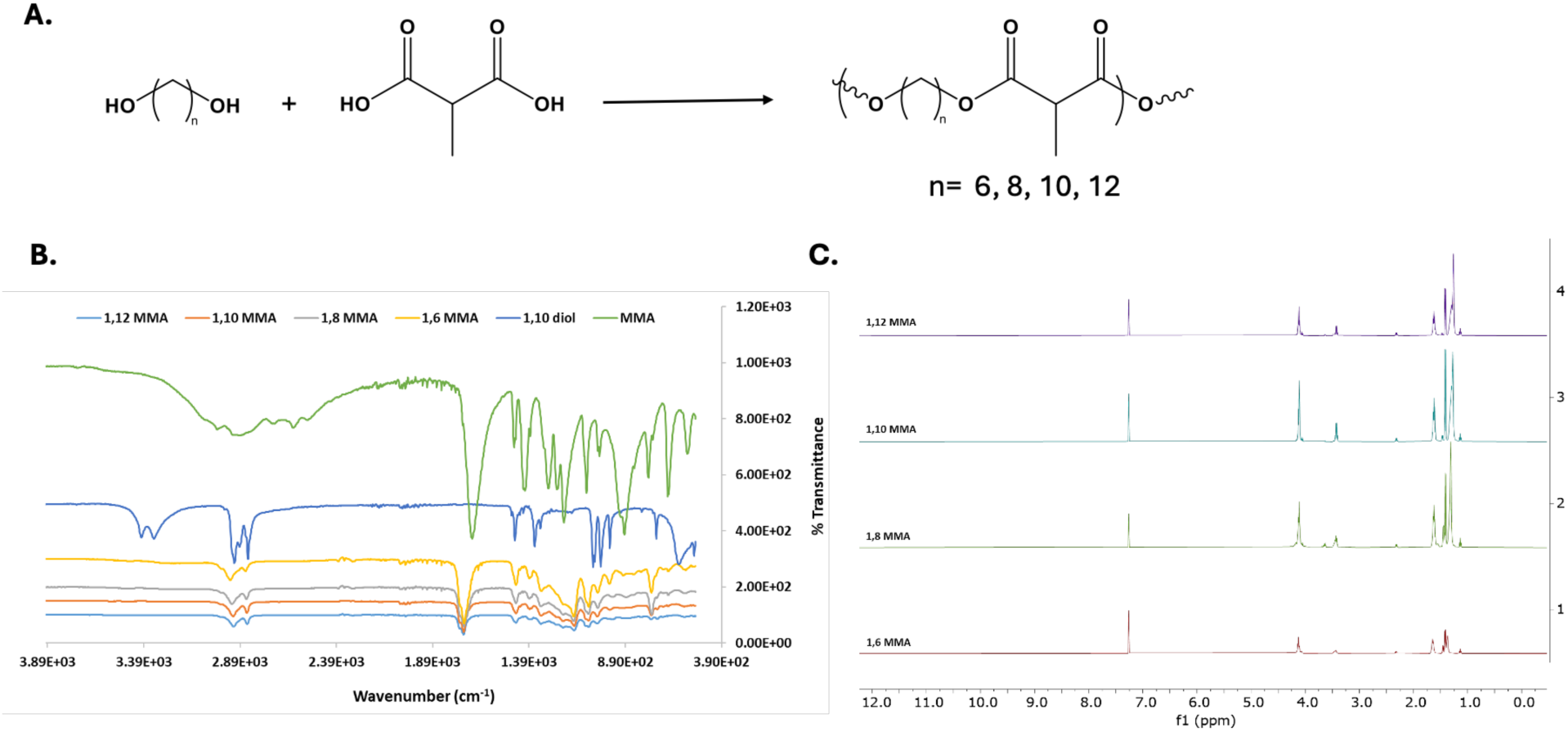
Synthesis and characterization of MMA polymer. A. Synthetic scheme of different chain length polymers of MMA. B. ATR-FTIR data showing the functional groups of alcohols, acids and esters (1735-1750 cm^-1^). C. Polymer generation was confirmed using 1H NMR, with characteristic peak of MMA **δ** = 4.13 ppm and 1,6 hexanediol **δ** = 3.45 ppm.

### 2.2 1,6 MMA has a higher immunosuppressive potential in inhibiting T cell activation

As mentioned before, MMA was previously reported to enhance CD8+ T cell exhaustion^23^. However, the concentrations used in previous studies suggested that free MMA would get rapidly metabolized and would not have long lasting effects. Therefore, we initially wanted to understand effect of this novel synthesized MMA polymer on T cell activation. CD3+ T cells were isolated from the spleen and activated in-vitro with plate bound αCD3 and soluble αCD28. We treated the cells with different doses of 1,6 MMA for 24 hours. **Fig S2A** demonstrates the gating strategy used to analyze these cells after flow cytometry. Initially we wanted to understand whether there is a change in CD25 (an early T cell activation marker and receptor component for IL2) expression in activated T cells treated with either soluble MMA (Free MMA) and 1,6 MMA. The data suggested that 1,6 MMA was more effective in causing a significant downregulation of CD25 frequency in both CD4+ and CD8+ T cells (**Fig S3A and B**). We next wanted to understand what happens to the IL2 secreted by these cell types. We observed that both Free MMA and Free diol, the monomers used to generate 1,6 MMA, could reduce IL2 secretion. However, the polymer was more robust in causing this phenomenon (**Fig S3C**). Overall, the data suggested that 1,6 MMA was more effective form of therapy in reducing pro-inflammatory T cell responses.

### 2.3 1,6 MMA reduces T cell activation by causing dysregulation in the IL2-pSTAT5 signaling axis

Our initial observations suggested that 250ug/ml of 1,6 MMA was effective in reducing CD25 frequency in T cells and cause effective reduction in IL2 levels. We wanted to further understand the mechanism behind this. We initially asked the question, whether CD25 expression also reduces or whether there is only reduction of overall CD25 frequency? **Fig 2A and B suggests** that 1,6 MMA causes significant reduction in CD25 MFI levels. Next, we wanted to understand whether there is a dose dependent effect of 1,6 MMA. Interestingly we observed that there was a dual role in this polymer. Low concentrations of 1,6 MMA (12.5ug/ml-50ug/ml) could cause significant reduction in CD25 percentage and expression. However, in both CD4+ and CD8+ T cells, CD25 frequency and expression was upregulated when cells were treated with concentrations of 62.5 and 125ug/ml (**Fig S4A-D**). As mentioned before 250ug/ml (concentration of interest) was the most effective in causing CD25 downregulation. However, the IL2 reduction due to 1,6 MMA happened in a dose dependent manner (**Fig S4I and S3C**). This suggested that although IL2 and CD25 expression are linked, there could be a delicate balance of IL2 dependent signaling required in causing significant CD25 downregulation.

**Figure 2:**
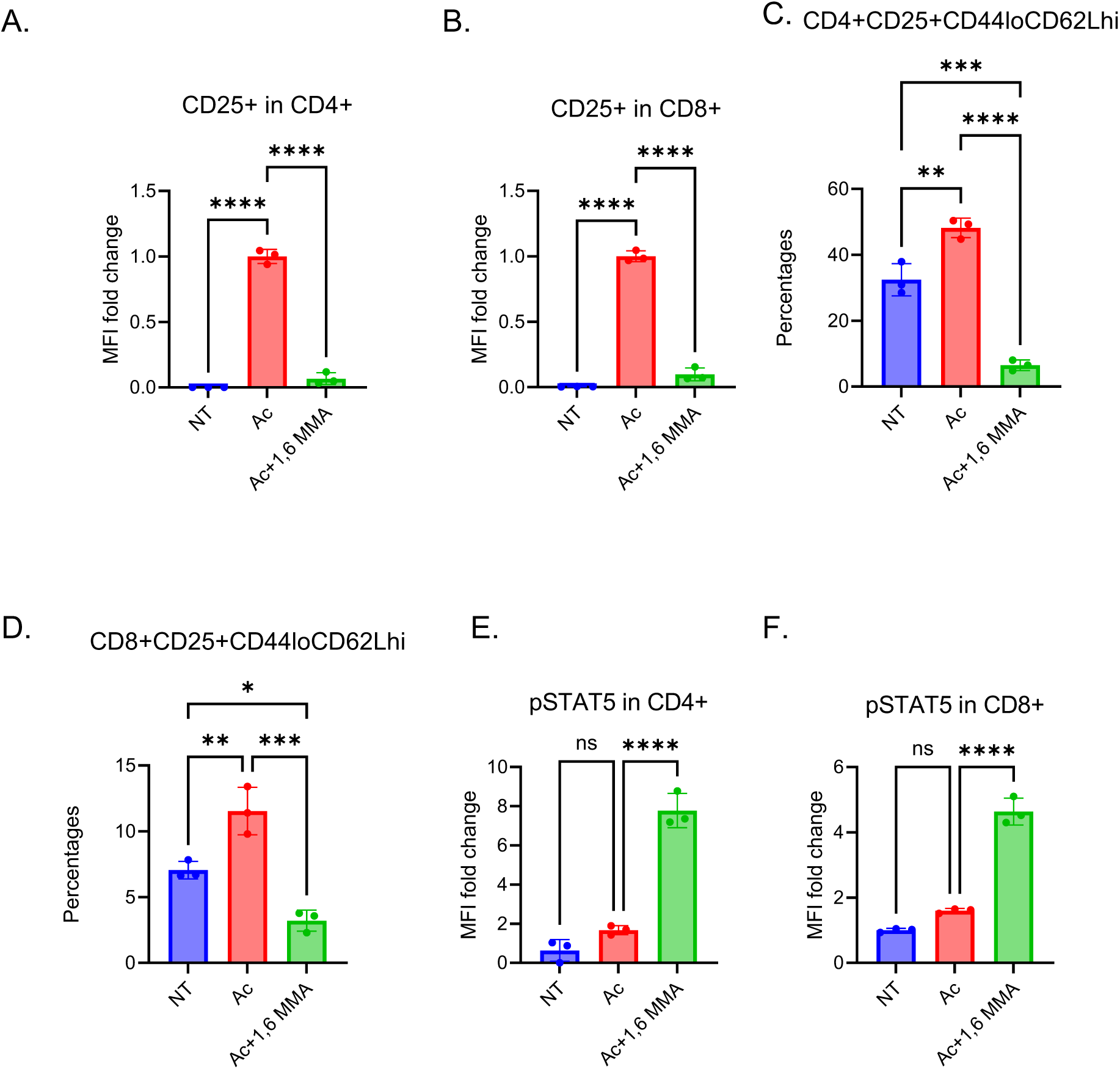
1,6 MMA inhibits T cell activation by an IL2 dependent mechanism. Quantification for flow cytometry analysis to understand the effect of 1,6 MMA ( 250µg/ml) on in-vitro activated T cells was performed. MFI fold change of A. CD25 of CD4+ T cells, B. CD25 of CD8+ T cells were quantified. Frequencies of C. CD4+CD25+CD44loCD62Lhi and D. CD8+CD25+CD44loCD62Lhi were calculated. Next, MFI fold change of pSTAT5 in E. CD4+ and F. CD8+ T cells were quantified. Student’s t test (A-C) and One-way Anova (D-F) were used to calculate statistical differences where *, **, *** and **** are for p values less than 0.05, 0.01, 0.001 and 0.0001 respectively.

We next wanted to confirm whether this loss in IL2 and CD25 causes loss of particular T cell populations. We stratified the CD25 expressing cells using CD44 and CD62L to understand what happens to the memory T cell formation with 1,6 MMA treatment. 24 hours being a short time point, we did not expect 1,6 MMA to have a profound impact on the T cell effector memory compartment. We therefore looked at the frequencies of the T cell population, CD4+CD25+CD44loCD62Lhi and CD8+CD25+CD44loCD62Lhi where we observed a significant loss in the former population. These populations could possibly be the regulatory T cell (Treg) compartment. This makes sense as Tregs require IL2 for survival and function^24^. This suggested that 1,6 MMA causes pan-T cell IL2 deprivation with activation. (**Fig 2C, 2D, S4E and S4F**).

Since pSTAT5 transcription factor plays an important role in regulating the IL2 signaling pathway in T cells^25^, we analyzed the intracellular amounts of pSTAT5 levels in both CD4+ and CD8+ T cells. Interestingly, we observed that pSTAT5 levels were abnormally high in the T cell compartments treated with 1,6 MMA (250µg/ml) (**Fig 2E and F**). This suggested that very high pSTAT5 amounts was negatively regulating IL2 secretion and CD25 expression. Finally, pSTAT5 levels increased in T cells with 1,6 MMA treatment in a dose dependent manner (**Fig S4G and H**)

### 2.4 1,6 MMA enhances cell death by increasing apoptosis in activated T cells

Since 1,6 MMA had an immunosuppressive effect, we wanted to confirm it was not due to non-specific toxic effects of the polymer. We initially used flow cytometry to calculate the frequency of viable T cells with polymer treatment at different doses. 1,6 MMA only increased cell death in activated T cells with significant increase in frequencies of cells positive for the viability dye. The increase in this percentage occurred in a dose dependent manner (**Fig 3A, 3B**, **S5A-D**).

**Figure 3:**
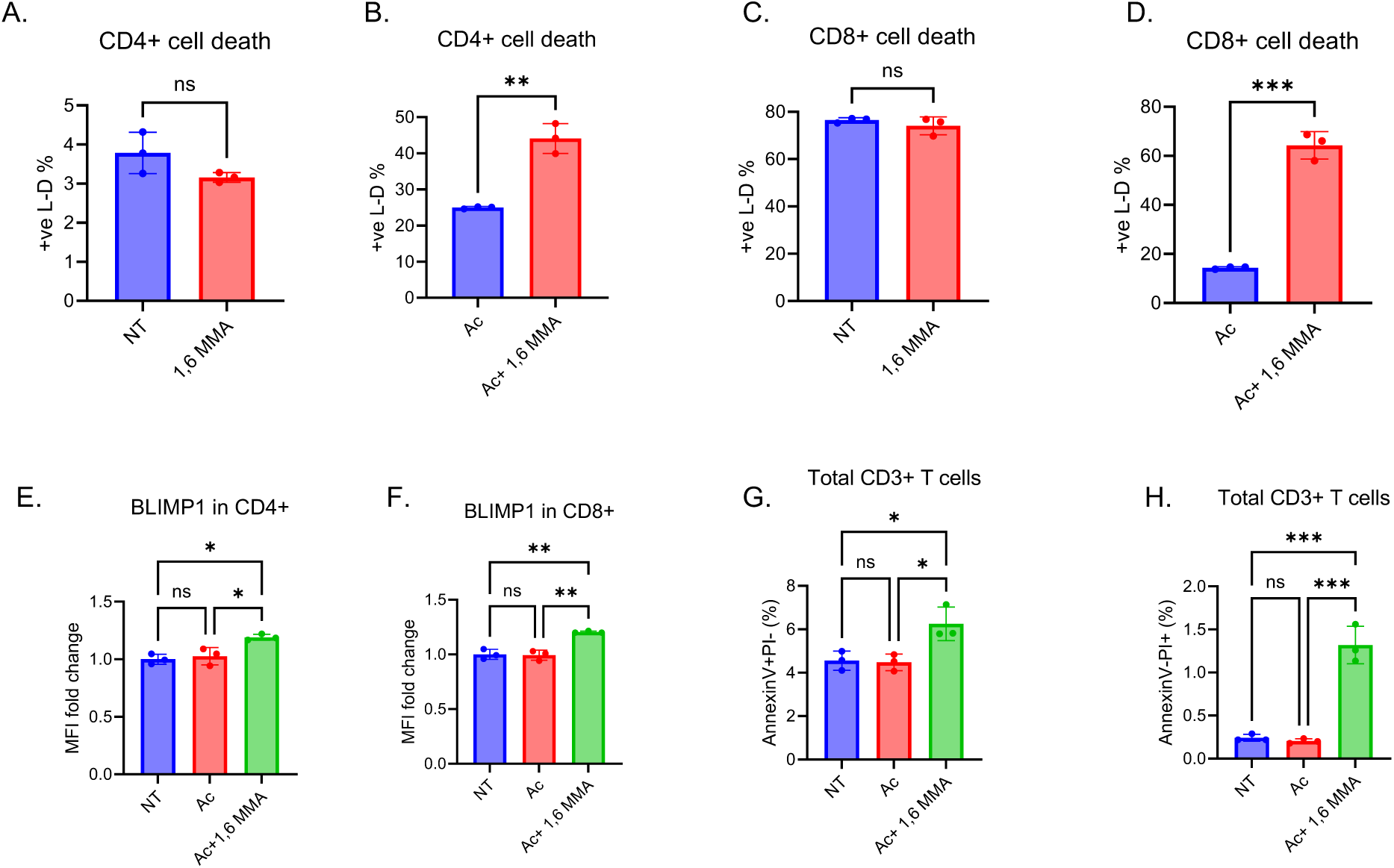
1,6 MMA inhibits T cell responses by enhancing apoptosis in activated cells. Quantification for flow cytometry data was performed. Bar graphs showing frequency of cells positive for viability dye (cell death) between control and 1,6 MMA (250µg/ml) treatment in A. un-activated CD4+, B. activated CD4+, C. un-activated CD8+ and D. activated CD8+ T cells. Next, graphs for calculating BLIMP-1 MFI fold changes were constructed for E. CD4+ and F. CD8+ T cells. Finally, specific flow analysis was performed using PI and Annexin V-FITC stains to calculate frequencies of G. AnnexinV+PI-(early apoptosis) and H. AnnexinV-PI+ (necrosis). Student’s t test (A-D) and One way Anova (E-H) were used to calculate statistical differences where *, **, *** and **** are for p values less than 0.05, 0.01, 0.001 and 0.0001 respectively.

This increase in cell death made us wonder whether the polymer has preference over a particular form of cell death pathway. We initially tested BLIMP-1, an intracellular protein, involved in apoptosis^26^. Additionally, pSTAT5 enhances BLIMP-1 expression which negatively regulates IL2 responses in T cells^27,28^. We observed an increase in BLIMP-1 expression in activated T cells which was dependent on the increasing doses of the polymer treatment (**Fig 3E, F and Fig S5E,F**). For further confirmation, we performed PI vs Annexin V flow experiments which further confirmed out hypothesis that suggested that 1,6 MMA enhanced early apoptosis (AnnexinV+PI-) and necrosis (AnnexinV-PI+) in activated T cells treated with the polymer (**Fig 3G, 3H, S5G, S5H**). To ensure this effect is specific for activated T cells, we analyzed early apoptosis and necrosis in CD4+ T cells, CD8+ T cells, CD11b+ (macrophage) and CD11c+ (dendritic) cells without activation. 1,6 MMA polymer at increasing doses did not causes significant changes in cell death (**Fig S6**) in these cell types, confirming the fact that the depletion of activated T cells occurs due to increased apoptosis/necrosis with a loss in IL2 signaling.

### 2.5 1,6 MMA suppresses T cell activation by enhancing oxidative phosphorylation and reactive oxygen species

Since metabolism plays a major role in determining the activation state of T cells, we performed seahorse experiments to understand the basal and maximal respiration of activated CD3+ T cells when treated with different doses of 1,6 MMA. The Oxygen consumption rate (OCR) curves from our seahorse analysis suggested an increase in oxidative phosphorylation when activated T cells were treated with different doses of 1,6 MMA (**Fig S7A**). We then quantified the basal and maximal respiration to understand efficiency of this metabolic pathway. Both basal and maximal respiration increased with 1,6 MMA treatment apart from 250ug/ml which does not cause much change in basal respiration (**Fig 4A, 4B, S7B and S7C**).

**Figure 4:**
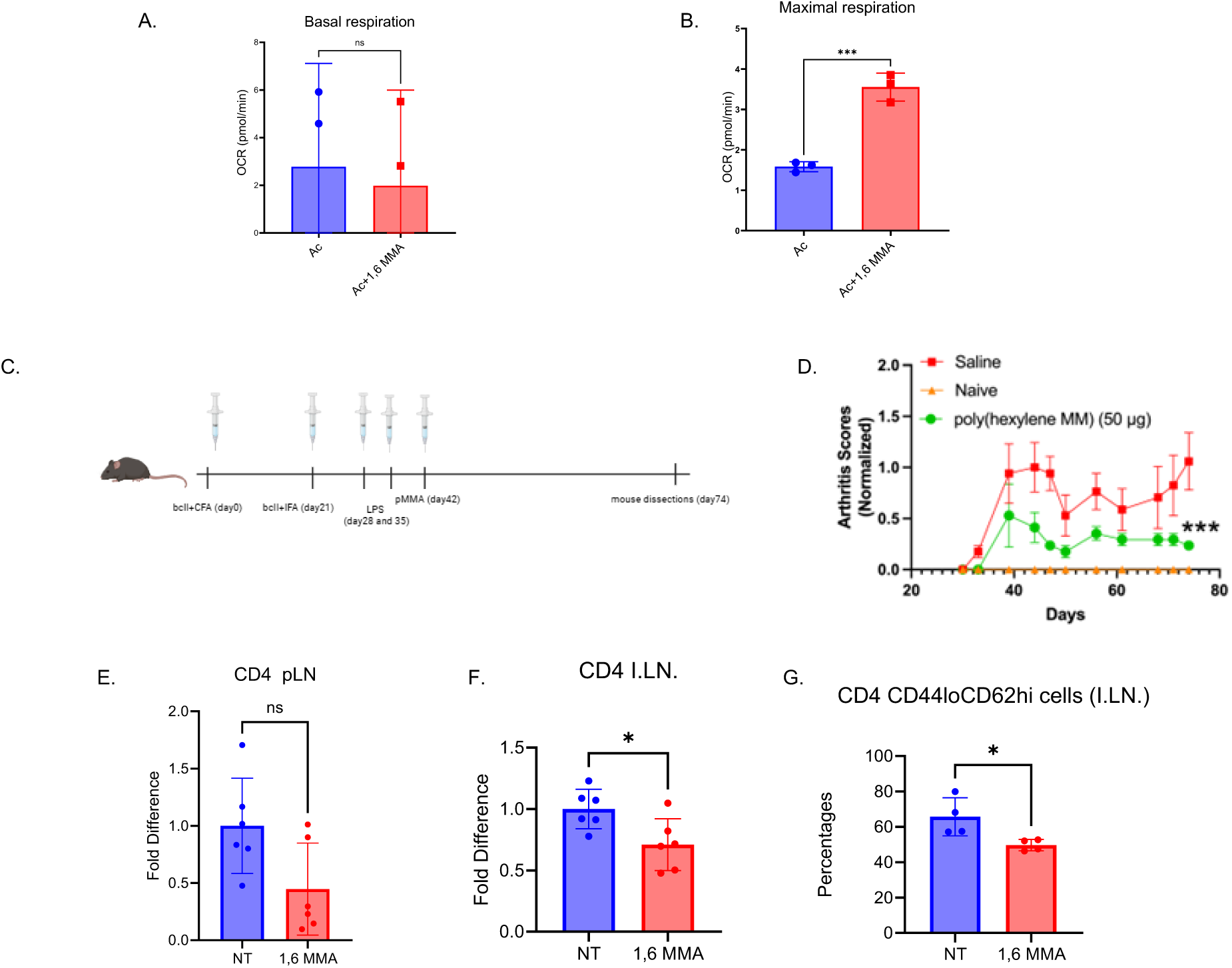
1,6 MMA enhances OCR in activated T cells and reduces disease symptoms in CIA mice. Seahorse analysis was done to calculate A. basal respiration and B. maximal respiration in T cells where we compare activated controls with 1,6 MMA (250µg/ml) treatment. C. experimental strategy showing the 1,6 MMA treatment regime and induction of CIA in mice. D. Arthritis scores calculate for naïve, CIA control (saline) and 1,6 MMA treated mice. Flow analysis was done and frequency of CD4+ T cells were calculated for E. popliteal lymph nodes (pLN) and F. inguinal lymph nodes (I.LN.) in CIA controls (saline/NT) and treatment mice (1,6 MMA). G. Frequency of CD4+CD44loCD62Lhi cells in I.LN. were calculated. Student’s t test was used to calculate statistical differences where *, **, *** and **** are for p values less than 0.05, 0.01, 0.001 and 0.0001 respectively.

Oxidative phosphorylation is linked to the electron transport chain in mitochondrial respiration which eventually causes the generation of superoxide (O_2_^-^)^29^. This contributes to the formation of reactive oxygen species which is known to both enhance and suppress immune responses depending on the intracellular amounts^30,31^. We performed flow cytometry analysis and observed increase in ROS levels with increasing doses of 1,6 MMA used to treat activated T cells (**Fig S7D**). This suggested that increase in oxidative phosphorylation caused inhibition of T cell activation by enhancing ROS production.

### 2.6 1,6 MMA suppresses disease in mouse models of rheumatoid arthritis by a T cell dependent mechanism

Our previous observations related to the invitro studies show us that 1,6 MMA specifically inhibits activated T cells, reduces IL2 production and enhances apoptosis. We therefore wanted to understand whether this polymer can reduce inflammation by reducing T cell responses in mice models of Rheumatoid Arthritis. We therefore used two separate mouse models to confirm our claim. First, we used an antigen driven model system using bovine collagen to elicit inflammation mice using CFA or IFA. 1 week post LPS treatment we inject a single dose of 1,6 MMA (50µg/mouse) via the intraperitoneal route (**Fig 4C**). We observed a significant drop in disease scores in mice with regards to paw and digit inflammation (**Fig 4D**). We wanted to confirm the immunological impact the polymer had on these mice. We analyzed the T cell profile from the draining lymph nodes which are the popliteal and inguinal lymph nodes. There was a modest but not significant decrease in CD4+ T cells in popliteal lymph nodes suggesting the possibility of depletion of activated T cells (**Fig 4E**). However, in the inguinal lymph nodes there was a significant loss of CD4+ T cells (**Fig 4F**). This would eventually reduce T cell dependent proinflammatory response causing the observed differences in disease scores. Interestingly, we observed a decrease in the CD44loCD62Lhi CD4 T cells (possibly regulatory T cells) which require IL2 for survival (**Fig 4G**). This observation was consistent to our in vitro studies suggesting the possibility of the same IL2 dependent in-vivo mechanism used by 1,6 MMA to reduce chronic inflammation.

### 2.7 1,6 MMA reduces human CD3+ T cell activation via a similar IL2 dependent mechanism

Following the mouse in vivo analysis, we investigated the impact of different concentrations of 1,6 MMA (labelled pMMA) on isolated human CD3⁺ T cells. Experimental groups included different concentrations of Ac+1,6 MMA and Ac+MMA, with non-treated (NT) and activated (Ac) samples used as controls. **Fig S8** shows that 1,6 MMA showed no toxic effects on human CD3⁺ T cells, even at the highest tested concentration (250 µg/mL). **Fig 5 A, 5B, S9F and S9G** showed that 1,6 MMA treatment reduced CD25 expression significantly in both CD8⁺ and CD4⁺ T cells compared to the activated controls. Notably, CD25 expression levels following 1,6 MMA treatment returned to a homeostatic range comparable to NT controls, suggesting a normalization of T-cell activation. **Fig 5C** further confirmed the reduced IL-2 expression following 1,6 MMA treatment relative to the activated control. **Fig S9H** depicted a similar increase in BLIMP-1 expression when CD8+ T cells were treated with 250μg/ml of 1,6 MMA. Unlike the mice experiments we did not observe a significant change in pSTAT5 levels in human CD3+ T cells (**Fig S9I and K**). Collectively, this indicated an overall reduction in T-cell activation upon 1,6 MMA treatment.

**Figure 5:**
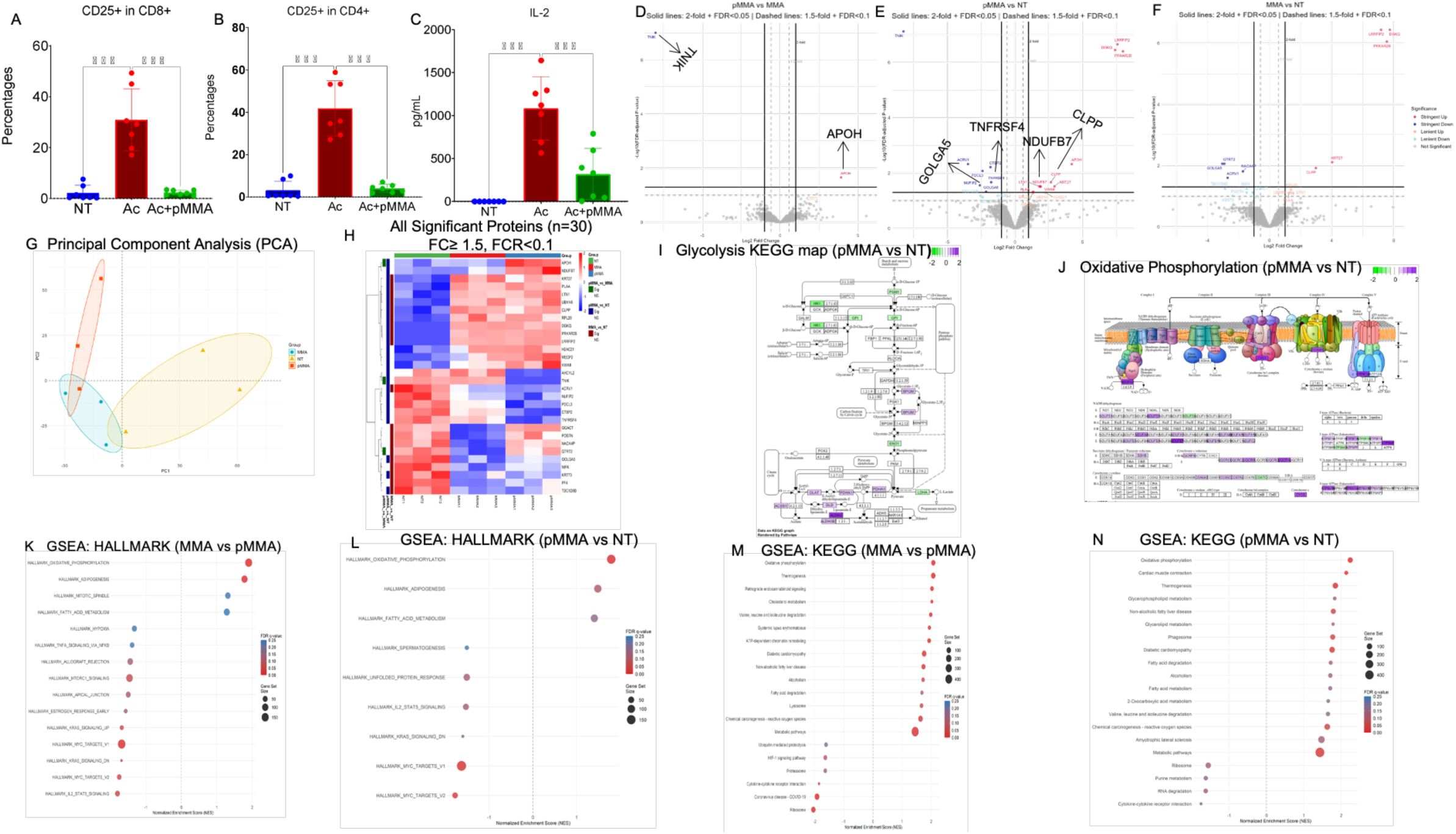
Unsupervised proteomic analysis reveal metabolic and functional changes in pMMA and MMA compared to NT (Ac) A. Flow cytometric quantification of CD25⁺ T cells within the CD8⁺ and B. CD4⁺ T cells under the indicated conditions (NT, Ac, Ac+pMMA). Individual data points represent biological replicates; bars indicate mean ± SEM. Statistical significance is indicated above bars (statistical test as described in Methods). C. Secreted IL-2 levels measured by ELISA. D. Differential protein abundance for pMMA vs MMA, E. pMMA vs NT, and F. MMA vs NT. The x-axis shows log_2_-fold change and the y-axis shows −log_10_(FDR). Proteins are classified into five significance categories: *stringent upregulated* or *downregulated* (solid lines; FDR < 0.05 and |log_2_FC| ≥ 1), *lenient upregulated* or *downregulated* (dashed lines; FDR < 0.10 and |log_2_FC| ≥ log_2_1.5), and *not significant*. Proteins of particular biological interest are pointed out and labeled to facilitate interpretation. G. Principal component analysis (PCA) of normalized proteomic profiles demonstrates clear separation between NT, MMA, and pMMA samples, indicating global proteomic reprogramming associated with treatment condition. Ellipses denote group clustering. H. Heatmap of significantly altered proteins (n = 30) across all pairwise comparisons (|fold change| ≥ 1.5, FDR < 0.1). Protein abundances are shown as row-wise Z-scores to emphasize relative expression patterns across samples. KEGG pathway maps of I. Glycolysis/Gluconeogenesis and J. Oxidative Phosphorylation in pMMA vs NT. Proteins are colored according to log_2_ fold change, highlighting pathway-level metabolic remodeling associated with the pMMA condition. Gene set enrichment analysis (GSEA) of proteomic changes using Hallmark gene sets for K. MMA vs pMMA and L. pMMA vs NT, and KEGG gene sets for M. MMA vs pMMA and N. pMMA vs NT. Dot plots display normalized enrichment scores (NES), with dot size representing gene set size and color indicating FDR-adjusted significance. One way Anova (A-C) were used to calculate statistical differences where *, **, *** and **** are for p values less than 0.05, 0.01, 0.001 and 0.0001 respectively.

### 2.8 1,6 MMA causes significant proteomic difference in activated human CD3+ T cells

Following flow cytometry analysis, we wanted to further characterize protein-level alterations induced by 1,6 MMA compared with MMA (free/soluble MMA) and NT controls. To this end, we performed a label-free relative protein abundance analysis across the three conditions. In **Fig 5G**, principal component analysis (PCA) was conducted on log2-transformed and normalized protein intensities to assess global expression patterns. PCA revealed that PC1 and PC2 accounted for 24.5% and 19.4% of the total variance, respectively. 1,6 MMA and NT samples showed clear separation, whereas MMA samples partially overlapped with both NT and 1,6 MMA, indicating moderate clustering overlap. The distinct separation between 1,6 MMA and NT reflects a pronounced proteomic shift following 1,6 MMA treatment, while the overlap observed with MMA suggests a less distinctive effect. No strong outliers were detected, indicating consistent data quality across samples. **Fig S10** also shows a bell-curve distribution of log2-transformed protein intensities across all samples, suggesting proper normalization^32^.

**Fig 5D-F** show volcano plots depicting differential protein expression between 1,6 MMA, MMA, and NT conditions. Significantly up- or downregulated proteins were highlighted, while non-significant proteins were shown in gray. **Fig 5D** compares 1,6 MMA and MMA samples where we identified a limited number of significantly altered proteins. One of the significant hits was TNIK. It was downregulated in 1,6 MMA compared with MMA. Previous studies have shown that TNIK-deficient CD8⁺ T cells exhibit increased apoptosis and impaired recall responses^33^. **Fig 5E** also showed upregulation of proteins such as NDUFB7, VWA8, and CLPP, and downregulation of TNFRSF4 and GOLGA5 between 1,6 MMA and NT samples. NDUFB7 contributed to mitochondrial complex I assembly and stability^34^, while CLPP supports sustained oxidative phosphorylation^35^. Conversely, downregulation of TNFRSF4 may contribute to the suppression of pathogenic T-cell activation^36^. **Fig 5H and S11** presents heatmaps of all significantly differentially expressed proteins (n = 30; fold change ≥ 1.5, FDR < 0.1) across NT, MMA, and 1,6 MMA conditions. Among the most prominently upregulated proteins in 1,6 MMA relative to both NT and MMA was APOH (β2-glycoprotein I). APOH has often been linked to autoimmune disease progression^37^. Conversely, 1,6 MMA treatment resulted in downregulation of TNF receptor– and NF-κB–related signaling pathways, highlighted by a significant reduction in TNFRSF4 (OX40) expression.

**Fig 5I and J** signifies KEGG pathway maps comparing 1,6 MMA versus NT in glycolysis/gluconeogenesis and oxidative phosphorylation pathways, respectively. As shown in **Fig I,** there was a marked downregulation of protein associated with reduced glycolytic activity, alongside upregulation of proteins associated with increased flux through the TCA cycle. **Fig 5J** demonstrated selective remodelling of mitochondrial oxidative phosphorylation in 1,6MMA-treated cells, characterized by increased expression of subunits across complexes I, III, IV, and V. Upregulation of complex I supports NADH oxidation and reactive oxygen species generation. Several complex III subunits, including UQCRC, UQCRH, and CYC1, were also upregulated. As complex III is a major site of mitochondrial ROS production, this suggests increased electron flux and superoxide generation^38^.

**Fig 5K and L** present GSEA hallmark analyses. **Fig 5K** shows reduced Hallmark IL-2–STAT5 signalling in 1,6 MMA compared with MMA, as well as increased oxidative phosphorylation in MMA relative to 1,6 MMA. **Fig 5L** shows increased oxidative phosphorylation in 1,6 MMA compared with NT. **Fig 5M and N** display GSEA-KEGG analyses, confirming these findings. Overall, this data further reconfirmed the roles of 1,6 MMA in dysregulation of mitochondrial function, increase in ROS levels and IL2 dependent T cell suppression.

## 3. Discussion

In this study we describe a novel metabolite-based polymer that suppresses both innate and adaptive immune responses and thereby reverse symptoms of rheumatoid arthritis and inflammation. Targeting immunometabolism to influence immune responses and maintain immune homeostasis has garnered attraction as alternative therapeutic approaches. In that context, metabolites have been engineered in the form of nanoparticles (NPs) and polymers to achieve stable immunomodulation to improve disease symptoms. Recent work from the lab showed the efficiency of polyethylene succinate nanoparticles (PES NPs) in reducing M1 macrophage responses which in turn reduced T cell responses invitro. The study showed the importance of this drug when used in combination with anti-PD1 therapy to efficiently stop tumor growth in melanoma mouse models^20^. Other studies from the lab showed the development of alpha-ketoglutarate (aKG) in the polymer backbone (paKG microparticles) which was used as a delivery system for PFK15, a known glycolysis inhibitor. paKG by itself also enhances oxidative phosphorylation in immune cells. In combination, this formulation had the capability to reduce dendritic cell (DCs) mediated immune activation and reduce symptoms of RA in CIA mice^21,39, 40^. This opens new doors for therapeutic interventions to treat RA and improve existing treatments such as methotrexate.

Methyl malonic acid (MMA), being a small molecule metabolite and a byproduct of the propionate metabolism pathway, has the potential to be used for immunosuppression. MMA is generated during the conversion of D-methylmalonyl-CoA to L-Methylmalonyl-CoA which eventually gets converted to Succinyl-CoA. This end-product gets incorporated into the TCA cycle. Accumulation of MMA has been linked to a defect in the enzyme Methylmalonyl-CoA mutase which helps in the formation of the mentioned end-product. This accumulation causes mitochondrial dysfunction, cell damage and has been linked to inborn errors which could have severe consequences^41,42^. With regards to the tumor microenvironment, the propionate metabolism pathway remains dysfunctional, causing an accumulation of MMA which promotes tumor growth and reduces anti-tumor immune cell responses^43^. Specifically, it has been reported that MMA has the capability to enhance CD8+ T cell exhaustion and reduce T cell activation. MMA has been shown to increase immunosuppressive transcriptional reprogramming by increasing the expression of the exhaustive regulator TOX. The study also focused on mitochondrial dysfunction and abnormalities in the TCA cycle that played a contributing factor in reduced T cell responses^23^. Interestingly, this shows the potential of using this molecule to suppress uncontrolled immune responses.

To implement this idea we first wanted to cross the hurdle of stabilizing the exposure time of MMA when added to activated T cells. This idea stems from the fact that MMA is a hydrophilic small molecule and in homeostatic conditions could get metabolized quickly. Therefore, to observe effective differences in reducing RA inflammation we must use higher concentrations of MMA. This reduces the impact of using it as an alternative therapy and could have serious side effects. Biomaterials are excellent for delivery of metabolic pathway inhibitors and metabolites that themselves act as immune cell function modulators^20,21,39, 40, 44–47^. Post characterization of this polymer we wanted to understand whether the MMA concentrations are detectable for an effective period of time. Detectable amounts of the polymer were observed in mice blood even after 21 days of a single injection via the i.p. route (50µg /mouse). This confirmed the stability of the MMA release from the polymer in-vivo.

Our in-vitro T cell data showed us the capability of 1,6 MMA in reducing T cell activation by causing significant reduction in CD25 and IL2 expression as early as 24 hours post activation in both mice and human T cells. In fact, the polymer caused a robust drop in CD25 expression in both CD4+ and CD8+ T cells suggesting pan T cell suppression. Additionally, the MMA monomer was incapable of causing similar changes in T cell responses, reconfirming the advantage of the polymer as a delivery system. The loss of IL2 in both human and mice T cells piqued our interest in understanding whether the polymer had a direct influence in affecting the IL2 signaling pathway. Therefore, our observations of enhanced pSTAT5 accumulation in mouse T cells was surprising as it suggested that effective IL2 mediated signaling required a delicate balance in pSTAT5 amounts. The paradoxical nature of this observation suggested that high pSTAT5 accumulation could accelerate the expression of the negative regulators of IL2 signaling pathway. This motivated us to investigate the expression of BLIMP-1^27,28^ which was enhanced both in activated CD4+ and CD8+ T cells treated with 1,6 MMA suggesting that high pSTAT5 enhanced BLIMP-1 expression which causes a drop in IL2 production. We wanted to know whether this loss in T cell responses was due to non-specific toxicity. Our cell death studies confirmed that 1,6 MMA enhanced apoptosis only in activated T cells which makes sense as activated T cells require IL2 for their long-term survival. Our human T cell proteomics data also suggested that 1,6 MMA causes global protein level changes with significant changes in TNF and NFκB pathways that are involved in robust T cell activation^48,49^. This leads to the fact that 1,6 MMA has robust immunosuppressive properties

Metabolically, 1,6 MMA enhanced OCR amounts which in turn suggested a dysregulation in oxidative phosphorylation. Enhanced oxidative phosphorylation is known to reduce proinflammatory T cell responses^50^. This observation further solidifies the reason behind lowered IL2 production. Since oxidative phosphorylation is tied to superoxide formation, we wanted to understand the changes in ROS levels in T cells. 1,6 MMA caused significant increase in ROS levels in activated T cells which would further cause T cell anergy and apoptosis. Interestingly, increased ROS production causes the inactivation of phosphatases inside cells. This suggests that 1,6 MMA causes high accumulation of pSTAT5 via a ROS dependent mechanism. The mitochondrial dysfunction, changes in OXPHOS and ROS generation was surprisingly common in out human T cell proteomic dataset for the 1,6 MMA group. Finally, our in-vitro findings helped us understand the effect of 1,6 MMA in CIA mice. The reduction of disease symptoms was clearly due to a reduction in T cell responses as observed when we look at the CD4+ T cell loss in the draining lymph node compartment. Overall, this suggested that 1,6 MMA reduces disease symptoms of RA by inhibiting T cell responses by causing enhanced ROS production and disruption in IL2 dependent cell signaling. In conclusion, 1,6 MMA is a novel metabolite-based polymer that has the potential to be used as an alternative therapy to suppress uncontrolled T cell responses and reduce symptoms of rheumatoid arthritis in individuals.

## 4. Materials and Methods

### 4.1 Materials used in study

Methylmalonic acid (CAS: 516-05-2, Fisher Scientific) and 1,6-Hexanediol (CAS: 629-11-8) or 1,8-Octanediol (CAS: 629-41-4) or 1,10-Decanediol (CAS: 112-47-0) or 1,12-Dodecanediol (CAS: 5675-51-4) were purchased from Sigma.

### 4.2 Synthesis of polymer

The polymers were synthesized via a slightly modified thermal esterification protocol (Ref: Succinate based polymers drive immunometabolism in dendritic cells to generate cancer immunotherapy). Brief, an equimolar ratio (1:1) of Methylmalonic acid and a diol (1,6-Hexanediol, 1,8-Octanediol, 1,10-Decanediol or 1,12-Dodecanediol) was heated at 70 ^0^C under vacuum for 48 h. Following this, the reaction temperature was increased to 90 ^0^C for 1 h to drive the polymerization to completion. The resulting polymer were purified by dissolving the crude mixture in chloroform and precipitating them through dropwise addition into stirring methanol or water. The precipitate was collected via centrifugation (5000 rpm for 5 min at room temperature). Residual chloroform was removed using a rotary evaporator, and the polymers were further dried under vacuum for 48 h to ensure complete removal of solvent. The purified polymers were stored at 4 ^0^C prior to chemical characterizations and biological evaluations.

### 4.3 Cell culture

Mouse CD3+ primary T cells and human CD3+ T cells were cultured in RPMI (Gibco, USA) that was supplemented with 10% FBS (Atlas Biologicals, Mexico), 1% penicillin and streptomycin antibiotics (CORNING, USA) and incubated at 37^0^C having a 5% CO_2_ environment. For the T cell activation studies, cells were cultured in 96 well U bottom plates (Fisherbrand, USA) for 24 hours. T cells were seeded at a density of 100,000 cells per well. Cells were treated with different doses of 1,6 MMA (12.5µg/ml to 250µg/ml) and treated with a single dose of the monomers, MMA and 1,6 Hexane diol (125µg/ml) for 24 hours. After 24 hours cells were processed for flow cytometry-based cell analysis.

### 4.4 T cell isolation and activation

Mouse CD3+ T cells and Human CD3+ T cells were isolated from the spleen and PBMC respectively. To perform the isolation, a magnetic based separation, mouse CD3+ and human CD3+ T cell isolation kit was used (Biolegend, USA). 96 well U bottom plates were coated with either purified anti-mouse CD3+ or anti-human CD3+ antibodies (Biolegend, USA) at a concentration of 0.5µg/ml. Plate was incubated at 4^0^C for 12-18 hours. Post incubation, wells were washed once with 1X PBS (VWR, Radnor, PA). Human or Mouse CD3+ T cells were added to the coated wells at the mentioned seeding densities. Purified anti human/mouse CD28 antibody (Biolegend, USA) was added to the respective wells at a concentration of 0.5µg/ml. Cells were analyzed 24 hours post activation. For LC/MS analysis, human CD3+ T cells were activated with 10ng/ml of PMA (Thermoscientific Chemicals, USA) and 0.1 μM of Ionomycin (Thermoscientific Chemicals, USA)

### 4.5 Flow cytometry (table for flow antibodies)

Flow cytometry (FACS) staining buffer was prepared using a combination of 0.1% bovine serum albumin (VWR, Radnor, PA), 2 mM Na_2_EDTA (VWR, Radnor, PA) and 0.01% NaN_3_ (VWR, Radnor, PA). Intracellular staining were performed by using the eBioscience FoxP3/Transcription factor staining buffer set (Invitrogen, USA). Live/dead staining was conducted using fixable dye eF780 (Invitrogen, USA). For reactive oxygen species (ROS) analysis, CellROX Orange was used (Invitrogen, USA). All cell surface and intracellular antibodies were purchased and used as received. Antibodies were added to samples at specific concentrations post titration. Required compensation and FMOs were used as reference for data analysis. Novocyte Quanteon (Agilent) was utililized for cell analysis. For data analysis Flowjo software was used.

**Table 1.**
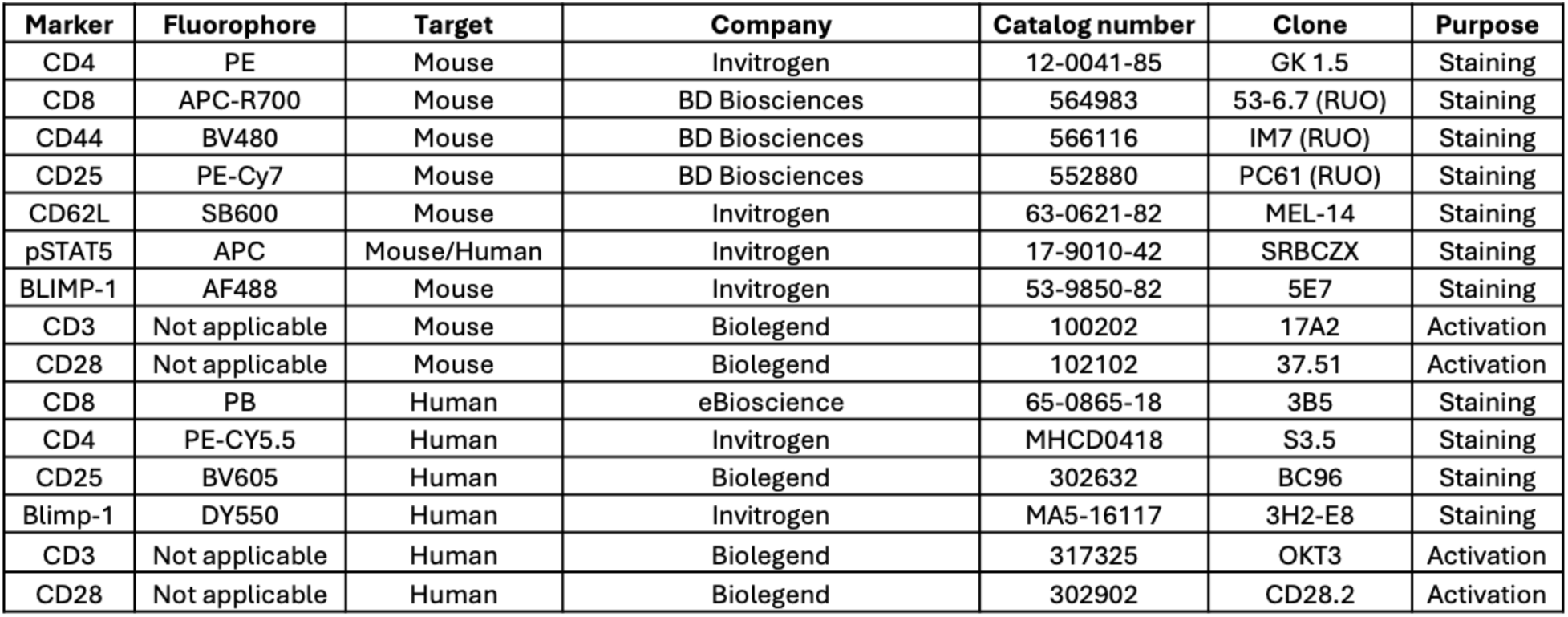
List of antibodies used for T cell activation and flow stainings.

### 4.6 ELISA

Human and Mouse CD3+ T cells were activated and treated with 1,6 MMA for 24 hours. Post activation cell supernatant was collected and IL2 ELISA was performed by using either mouse or human IL2 ELISA kit (Invitrogen, USA). Sample were analyzed for absorbance at 450nM using the BioTek Synergy H1 plate reader.

### 4.7 Seahorse analysis

Oxygen consumption rate (OCR) analysis was done on both human and mice CD3+ T cells. Seahorse protocol for OCR analysis was performed by processing cells using a previous established protocol from the lab^51^. Seahorse analysis was done using Seahorse Extracellular Flux XF-96) analyser (Seahorse Bioscience, North Billerica, MA).

### 4.8 In vivo experiments

The IACUC of Case Western Reserve University approved animal studies for rheumatoid arthritis). Eight-week-old female DBA/1j (Jackson Laboratories) mice were used to induce collagen induced arthritis (CIA) in mice. Previous protocol from the lab was followed for CIA induction using in mice^21,39,40^. 30 days post CIA induction mice were given a single dose of 1,6 MMA (50µg/mouse) via the intraperitoneal route. Mice were scored for both paw and digit inflammation at regular time intervals.

### 4.9 Sample preparation and protein extraction for human CD3+ T cells

Nine cell pellets were processed by the Lerner Research Institute’s Proteomics & Metabolomics Core for proteomic analysis by in-solution digestion followed by LC–MS/MS. Briefly, pellets were lysed in 6 M urea/Tris-HCl buffer, and protein concentration was assessed by BCA assay. Due to low protein abundance, samples were reduced with dithiothreitol and alkylated with iodoacetamide, after which the urea concentration was diluted with Tris-HCl. Proteins were digested overnight at 37 °C using trypsin (1:100, trypsin:protein). Digestion was quenched with trifluoroacetic acid, and peptides were desalted using C18 spin columns, dried by SpeedVac, and reconstituted in 0.1% formic acid prior to analysis.

Peptides were analyzed on a Bruker timsTOF Pro 2 mass spectrometer coupled to nanoLC using a C18 reversed-phase column and a 50-min acetonitrile gradient. Data were acquired in positive ion mode using data-dependent acquisition with PASEF for peptide fragmentation. Raw data were searched against the human SwissProtKB database (with common contaminants) using PEAKS Online, with trypsin specificity, carbamidomethylation as a fixed modification, oxidation of methionine as a variable modification, and a 1% false discovery rate at the peptide level. Proteins were reported with a minimum of two peptides, including at least one unique peptide.

### 4.10 Data processing and Statistical analysis

Raw files were processed using MaxQuant (v2.0.3.0) with label-free quantification (LFQ) against the human UniProt database.^52,53^. LFQ intensities were log_2_-transformed, and proteins detected in ≥70% of samples were retained. Variance stabilization normalization (VSN) was applied using the vsn package^54^, and proteins with CV >0.5 were excluded. Missing values were imputed using K-nearest neighbors (k=3). Quality control included PCA, distribution plots, and correlation analysis. Differential expression was analyzed using limma with empirical Bayes moderation^55^ for three comparisons: MMA vs. NT, 1,6 MMA vs. NT, and 1,6 MMA vs. MMA. Proteins were considered significant at |log_2_FC| ≥ 0.585 (1.5-fold) and FDR < 0.1.

### 4.11 Pathway Enrichment and Network Analysis

Gene set enrichment analysis (GSEA) was performed using clusterProfiler^56^ with proteins ranked by - log_10_(p-value) × sign(log_2_FC). Two databases were queried: KEGG^57^, and MSigDB Hallmark^58^. Parameters: minimum gene set = 10, maximum = 500, FDR < 0.25, 10,000 permutations.

### 4.12 Pathway Visualization

KEGG pathway maps were generated using the pathview R package^59^, overlaying protein fold changes onto pathway diagrams to visualize coordinated regulation.

### 4.13 Visualization

Figures were generated in R (v4.3.0) using ggplot2 and ComplexHeatmap^60^. Heatmaps display z-score normalized values with Ward’s hierarchical clustering. All figures were exported at 300 dpi in TIFF/PNG formats.

### 4.14 Software and Statistical Environment for LC/MS analysis

All proteomic analyses were performed in R (version 4.3.0) using Bioconductor (version 3.18). Key packages included: limma (v3.56.0), vsn (v3.68.0), clusterProfiler (v4.8.0), ComplexHeatmap (v2.16.0), and ggplot2 (v3.4.0). Complete analysis code is available at [repository link].

Statistical significance was set at p < 0.05 (two-tailed) unless otherwise specified. All data (Ritchie et al., 2015). This approach provides robust statistical inference even with small sample sizes by borrowing information across proteins to stabilize variance estimates. All data with respect to invitro mice and human T cell experiments along with invivo mice experiments were analyzed for significant statistical differences using One way Anova with Tukey’s post test. Student t-test was used when analyzing only two groups for statistical differences. P values <0.05 were considered statistically significant. Data analysis were performed using Microsoft Excel and GraphPad Prism 9.

## Acknowledgement

The authors would like to acknowledge grants NIH 1R01AI155907-01 and NIH R01AR078343.

## References

1. Smolen, J. S., Aletaha, D., & McInnes, I. B. (2016). Rheumatoid arthritis. Lancet (London, England), 388(10055), 2023–2038. 10.1016/S0140-6736(16)30173-8

2. Orozco, G., Rueda, B., & Martin, J. (2006). Genetic basis of rheumatoid arthritis. Biomedicine & pharmacotherapy = Biomedecine & pharmacotherapie, 60(10), 656–662. 10.1016/j.biopha.2006.09.003

3. Liao, K. P., Alfredsson, L., & Karlson, E. W. (2009). Environmental influences on risk for rheumatoid arthritis. Current opinion in rheumatology, 21(3), 279–283. 10.1097/BOR.0b013e32832a2e16

4. Malmstrøm, V., Trollmo, C., & Klareskog, L. (2004). The additive role of innate and adaptive immunity in the development of arthritis. The American journal of the medical sciences, 327(4), 196–201. 10.1097/00000441-200404000-00005

5. Scherger, S., Henao-Martinez, A. F., Franco-Paredes, C., Ngo, B. T., Grimshaw, A., Sah, R., Shah, S., Sillau, S., Bastias, A. G., Corbisiero, M. F., Kyllo, H. M., Stellern, J., & Shapiro, L. (2025). Circulating TNF-α levels in rheumatoid arthritis: a systematic review and meta-analysis and comparison to TNF-α levels in sepsis. Therapeutic advances in infectious disease, 12, 20499361251368006. 10.1177/20499361251368006

6. Pandolfi, F., Franza, L., Carusi, V., Altamura, S., Andriollo, G., & Nucera, E. (2020). Interleukin-6 in Rheumatoid Arthritis. International journal of molecular sciences, 21(15), 5238. 10.3390/ijms21155238

7. Dayer, J. M., Oliviero, F., & Punzi, L. (2017). A Brief History of IL-1 and IL-1 Ra in Rheumatology. Frontiers in pharmacology, 8, 293. 10.3389/fphar.2017.00293

8. Baslund, B., Tvede, N., Danneskiold-Samsoe, B., Larsson, P., Panayi, G., Petersen, J., Petersen, L. J., Beurskens, F. J., Schuurman, J., van de Winkel, J. G., Parren, P. W., Gracie, J. A., Jongbloed, S., Liew, F. Y., & McInnes, I. B. (2005). Targeting interleukin-15 in patients with rheumatoid arthritis: a proof-of-concept study. Arthritis and rheumatism, 52(9), 2686–2692. 10.1002/art.21249

9. Ammon, T., Zeiträg, J., Mayr, V., Benedicic, M., Holthoff, H. P., & Ungerer, M. (2025). Citrullinated Autoantigen-Specific T and B Lymphocytes in Rheumatoid Arthritis: Focus on Follicular T Helper Cells and Expansion by Coculture. ACR open rheumatology, 7(1), e11785. 10.1002/acr2.11785

10. Song, J., Schwenzer, A., Wong, A., Turcinov, S., Rims, C., Martinez, L. R., Arribas-Layton, D., Gerstner, C., Muir, V. S., Midwood, K. S., Malmström, V., James, E. A., & Buckner, J. H. (2021). Shared recognition of citrullinated tenascin-C peptides by T and B cells in rheumatoid arthritis. JCI insight, 6(5), e145217. 10.1172/jci.insight.145217

11. Higashioka, K., & Rao, D. A. (2024). DC-T cell power couples in rheumatoid arthritis joints. Immunity, 57(12), 2715–2717. 10.1016/j.immuni.2024.11.012

12. Luo, P., Wang, P., Xu, J., Hou, W., Xu, P., Xu, K., & Liu, L. (2022). Immunomodulatory role of T helper cells in rheumatoid arthritis : a comprehensive research review. Bone & joint research, 11(7), 426–438. 10.1302/2046-3758.117.BJR-2021-0594.R1

13. Toghi, M., Bitarafan, S., & Ghafouri-Fard, S. (2023). Pathogenic Th17 cells in autoimmunity with regard to rheumatoid arthritis. Pathology, research and practice, 250, 154818. 10.1016/j.prp.2023.154818

14. Solomon, D. H., Glynn, R. J., Karlson, E. W., Lu, F., Corrigan, C., Colls, J., Xu, C., MacFadyen, J., Barbhaiya, M., Berliner, N., Dellaripa, P. F., Everett, B. M., Pradhan, A. D., Hammond, S. P., Murray, M., Rao, D. A., Ritter, S. Y., Rutherford, A., Sparks, J. A., Stratton, J., … Ridker, P. M. (2020). Adverse Effects of Low-Dose Methotrexate: A Randomized Trial. Annals of internal medicine, 172(6), 369–380. 10.7326/M19-3369

15. Gogulescu, A., Blidisel, A., Soica, C., Mioc, A., Voicu, A., Jojic, A., Voicu, M., & Banciu, C. (2024). Neurological Side Effects of TNF-α Inhibitors Revisited: A Review of Case Reports. Medicina (Kaunas, Lithuania), 60(9), 1409. 10.3390/medicina60091409

16. Zuckerman, B. P., Gibson, M., Roy, R., Hughes, M., Mehta, D., Yang, Z., Adas, M., Ng, K., Russell, M. D., Cope, A., Norton, S., & Galloway, J. (2025). Abatacept and the risk of malignancy: a meta-analysis across disease indications. Rheumatology (Oxford, England), 64(6), 3280–3287. 10.1093/rheumatology/keaf114

17. Hu, T., Liu, C. H., Lei, M., Zeng, Q., Li, L., Tang, H., & Zhang, N. (2024). Metabolic regulation of the immune system in health and diseases: mechanisms and interventions. Signal transduction and targeted therapy, 9(1), 268. 10.1038/s41392-024-01954-6

18. Kornberg M. D. (2020). The immunologic Warburg effect: Evidence and therapeutic opportunities in autoimmunity. Wiley interdisciplinary reviews. Systems biology and medicine, 12(5), e1486. 10.1002/wsbm.1486

19. Zhang, S., Lv, K., Liu, Z., Zhao, R., & Li, F. (2024). Fatty acid metabolism of immune cells: a new target of tumour immunotherapy. Cell death discovery, 10(1), 39. 10.1038/s41420-024-01807-9

20. Jaggarapu, M. M. C. S., Pathak, S., Joseph, J. P., Swaminathan, S. J., Inamdar, S., Suresh, A. P., & Acharya, A. P. (2026). Delivery of Macrophage Activating Nanoparticles and Checkpoint Inhibitors Improves Melanoma Immunotherapy in Mice. Small (Weinheim an der Bergstrasse, Germany), 22(2), e10586. 10.1002/smll.202510586

21. Mangal, J. L., Inamdar, S., Suresh, A. P., Jaggarapu, M. M. C. S., Esrafili, A., Ng, N. D., & Acharya, A. P. (2022). Short term, low dose alpha-ketoglutarate based polymeric nanoparticles with methotrexate reverse rheumatoid arthritis symptoms in mice and modulate T helper cell responses. Biomaterials science, 10(23), 6688–6697. 10.1039/d2bm00415a

22. Wongkittichote, P., Cunningham, G., Summar, M. L., Pumbo, E., Forny, P., Baumgartner, M. R., & Chapman, K. A. (2019). Tricarboxylic acid cycle enzyme activities in a mouse model of methylmalonic aciduria. Molecular genetics and metabolism, 128(4), 444–451. 10.1016/j.ymgme.2019.10

23. ejero, J. D., Hesterberg, R. S., Drapela, S., Ilter, D., Raizada, D., Lazure, F., Kashfi, H., Liu, M., Silvane, L., Avram, D., Fernández-García, J., Asara, J. M., Fendt, S. M., Cleveland, J. L., & Gomes, A. P. (2025). Methylmalonic acid induces metabolic abnormalities and exhaustion in CD8^+^ T cells to suppress anti-tumor immunity. Oncogene, 44(2), 105–114. 10.1038/s41388-024-03191-1

24. Harris, F., Berdugo, Y. A., & Tree, T. (2023). IL-2-based approaches to Treg enhancement. Clinical and experimental immunology, 211(2), 149–163. 10.1093/cei/uxac105

25. Owen, D. L., & Farrar, M. A. (2017). STAT5 and CD4 ^+^ T Cell Immunity. F1000Research, 6, 32. 10.12688/f1000research.9838

26. Fu, S. H., Yeh, L. T., Chu, C. C., Yen, B. L., & Sytwu, H. K. (2017). New insights into Blimp-1 in T lymphocytes: a divergent regulator of cell destiny and effector function. Journal of biomedical science, 24(1), 49. 10.1186/s12929-017-0354-8

27. Nurieva, R. I., Podd, A., Chen, Y., Alekseev, A. M., Yu, M., Qi, X., Huang, H., Wen, R., Wang, J., Li, H. S., Watowich, S. S., Qi, H., Dong, C., & Wang, D. (2012). STAT5 protein negatively regulates T follicular helper (Tfh) cell generation and function. The Journal of biological chemistry, 287(14), 11234–11239. 10.1074/jbc.M111.324046

28. Roy, S., Ren, M., Li, P., Cui, K., Cao, Y., Fisk, B., Markowitz, T. E., Redekar, N., Sakamoto, K., Nagao, K., Oh, J., Spolski, R., Liao, W., Dubois, S. P., Kelsall, B. L., Zhao, K., Phelan, J. D., & Leonard, W. J. (2025). BLIMP1 negatively regulates IL-2 signaling in T cells. Science advances, 11(29), eadx8105. 10.1126/sciadv.adx8105

29. Yarosz, E. L., & Chang, C. H. (2018). The Role of Reactive Oxygen Species in Regulating T Cell-mediated Immunity and Disease. Immune network, 18(1), e14. 10.4110/in.2018.18.e14

30. Case, A. J., McGill, J. L., Tygrett, L. T., Shirasawa, T., Spitz, D. R., Waldschmidt, T. J., Legge, K. L., & Domann, F. E. (2011). Elevated mitochondrial superoxide disrupts normal T cell development, impairing adaptive immune responses to an influenza challenge. Free radical biology & medicine, 50(3), 448–458. 10.1016/j.freeradbiomed.2010.11.025

31. van der Veen, R. C., Dietlin, T. A., Karapetian, A., Holland, S. M., & Hofman, F. M. (2004). Extra-cellular superoxide promotes T cell expansion through inactivation of nitric oxide. Journal of neuroimmunology, 153(1-2), 183–189. 10.1016/j.jneuroim.2004.05.008

32. Rigotto-Caruso, G., Curtis, A., Kavanagh, K., & von Zeska Kress, M. R. (2025). Label-Free Quantitative Proteomic Analysis Reveals the Effects of Biogenic Silver Nanoparticles on *Fusarium keratoplasticum* and Their Therapeutic Potential in *Galleria mellonella* Larvae. ACS omega, 10(33), 37408–37418. 10.1021/acsomega.5c03275

33. Jaeger-Ruckstuhl, C. A., Hinterbrandner, M., Höpner, S., Correnti, C. E., Lüthi, U., Friedli, O., Freigang, S., Al Sayed, M. F., Bührer, E. D., Amrein, M. A., Schürch, C. M., Radpour, R., Riether, C., & Ochsenbein, A. F. (2020). TNIK signaling imprints CD8^+^ T cell memory formation early after priming. Nature communications, 11(1), 1632. 10.1038/s41467-020-15413-7

34. Panga, V., Kallor, A. A., Nair, A., Harshan, S., & Raghunathan, S. (2019). Mitochondrial dysfunction in rheumatoid arthritis: A comprehensive analysis by integrating gene expression, protein-protein interactions and gene ontology data. PloS one, 14(11), e0224632. 10.1371/journal.pone.0224632

35. Weyand, C. M., Wu, B., & Goronzy, J. J. (2020). The metabolic signature of T cells in rheumatoid arthritis. Current opinion in rheumatology, 32(2), 159–167. 10.1097/BOR.0000000000000683

36. Castro-Sánchez, P., Aguilar-Sopeña, O., Alegre-Gómez, S., Ramirez-Munoz, R., & Roda-Navarro, P. (2019). Regulation of CD4^+^ T Cell Signaling and Immunological Synapse by Protein Tyrosine Phosphatases: Molecular Mechanisms in Autoimmunity. Frontiers in immunology, 10, 1447. 10.3389/fimmu.2019.01447

37. Suresh, S., Demirci, F. Y., Jacobs, E., Kao, A. H., Rhew, E. Y., Sanghera, D. K., Selzer, F., Sutton-Tyrrell, K., McPherson, D., Bontempo, F. A., Kammerer, C. M., Ramsey-Goldman, R., Manzi, S., & Kamboh, M. I. (2009). Apolipoprotein H promoter polymorphisms in relation to lupus and lupus-related phenotypes. The Journal of rheumatology, 36(2), 315–322. 10.3899/jrheum.080482

38. O’Neill, L. A., Kishton, R. J., & Rathmell, J. (2016). A guide to immunometabolism for immunologists. Nature reviews. Immunology, 16(9), 553–565. 10.1038/nri.2016.70

39. Mangal, J. L., Inamdar, S., Le, T., Shi, X., Curtis, M., Gu, H., & Acharya, A. P. (2021). Inhibition of glycolysis in the presence of antigen generates suppressive antigen-specific responses and restrains rheumatoid arthritis in mice. Biomaterials, 277, 121079. 10.1016/j.biomaterials.2021.121079

40. Thumsi, A., Martínez, D., Swaminathan, S. J., Esrafili, A., Suresh, A. P., Jaggarapu, M. M. C., Lintecum, K., Halim, M., Mantri, S. V., Sleiman, Y., Appel, N., Gu, H., Curtis, M., Zuniga, C., & Acharya, A. P. (2025). Inverse-Vaccines for Rheumatoid Arthritis Re-establish Metabolic and Immunological Homeostasis in Joint Tissues. Advanced healthcare materials, 14(5), e2303995. 10.1002/adhm.202303995

41. Almási, T., Guey, L. T., Lukacs, C., Csetneki, K., Vokó, Z., & Zelei, T. (2019). Systematic literature review and meta-analysis on the epidemiology of methylmalonic acidemia (MMA) with a focus on MMA caused by methylmalonyl-CoA mutase (mut) deficiency. Orphanet journal of rare diseases, 14(1), 84. 10.1186/s13023-019-1063-z

42. Chen, T., Gao, Y., Zhang, S., Wang, Y., Sui, C., & Yang, L. (2023). Methylmalonic acidemia: Neurodevelopment and neuroimaging. Frontiers in neuroscience, 17, 1110942.10.3389/fnins.2023.1110942

43. Gomes, A. P., Ilter, D., Low, V., Drapela, S., Schild, T., Mullarky, E., Han, J., Elia, I., Broekaert, D., Rosenzweig, A., Nagiec, M., Nunes, J. B., Schaffer, B. E., Mutvei, A. P., Asara, J. M., Cantley, L. C., Fendt, S. M., & Blenis, J. (2022). Altered propionate metabolism contributes to tumour progression and aggressiveness. Nature metabolism, 4(4), 435–443. 10.1038/s42255-022-00553-5

44. Mangal, J. L., Inamdar, S., Yang, Y., Dutta, S., Wankhede, M., Shi, X., Gu, H., Green, M., Rege, K., Curtis, M., & Acharya, A. P. (2020). Metabolite releasing polymers control dendritic cell function by modulating their energy metabolism. Journal of materials chemistry. B, 8(24), 5195–5203. 10.1039/d0tb00790k

45. Thumsi, A., Swaminathan, S. J., Mangal, J. L., Suresh, A. P., & Acharya, A. P. (2023). Vaccines prevent reinduction of rheumatoid arthritis symptoms in collagen-induced arthritis mouse model. Drug delivery and translational research, 13(7), 1925–1935. 10.1007/s13346-023-01333-8

46. Mohan Chandra Sekhar Jaggarapu, M., Thumsi, A., Nile, R., D Ridenour, B., Khodaei, T., P Suresh, A., Esrafili, A., Jin, K., & P Acharya, A. (2023). Orally delivered 2D covalent organic frameworks releasing kynurenine generate anti-inflammatory T cell responses in collagen induced arthritis mouse model. Biomaterials, 300, 122204. 10.1016/j.biomaterials.2023.122204

47. Inamdar, S., Tylek, T., Thumsi, A., Suresh, A. P., Jaggarapu, M. M. C. S., Halim, M., Mantri, S., Esrafili, A., Ng, N. D., Schmitzer, E., Lintecum, K., de Ávila, C., Fryer, J. D., Xu, Y., Spiller, K. L., & Acharya, A. P. (2023). Biomaterial mediated simultaneous delivery of spermine and alpha ketoglutarate modulate metabolism and innate immune cell phenotype in sepsis mouse models. Biomaterials, 293, 121973. 10.1016/j.biomaterials.2022.121973

48. Mehta, A. K., Gracias, D. T., & Croft, M. (2018). TNF activity and T cells. Cytokine, 101, 14–18. 10.1016/j.cyto.2016.08.003

49. Daniels, M. A., & Teixeiro, E. (2025). The NF-κB signaling network in the life of T cells. Frontiers in immunology, 16, 1559494. 10.3389/fimmu.2025.1559494

50. Shi, Y., Zhang, H., & Miao, C. (2025). Metabolic reprogram and T cell differentiation in inflammation: current evidence and future perspectives. Cell death discovery, 11(1), 123. 10.1038/s41420-025-02403-1

51. Inamdar, S., Suresh, A. P., Mangal, J. L., Ng, N. D., Sundem, A., Wu, C., Lintecum, K., Thumsi, A., Khodaei, T., Halim, M., Appel, N., Jaggarapu, M. M. C. S., Esrafili, A., Yaron, J. R., Curtis, M., & Acharya, A. P. (2023). Rescue of dendritic cells from glycolysis inhibition improves cancer immunotherapy in mice. Nature communications, 14(1), 5333. 10.1038/s41467-023-41016-z

52. Cox, J., Hein, M. Y., Luber, C. A., Paron, I., Nagaraj, N., & Mann, M. (2014). Accurate proteome-wide label-free quantification by delayed normalization and maximal peptide ratio extraction, termed MaxLFQ. Molecular & cellular proteomics : MCP, 13(9), 2513–2526. 10.1074/mcp.M113.031591

53. Tyanova, S., Temu, T., & Cox, J. (2016). The MaxQuant computational platform for mass spectrometry-based shotgun proteomics. Nature protocols, 11(12), 2301–2319. 10.1038/nprot.2016.136

54. Huber, W., von Heydebreck, A., Sültmann, H., Poustka, A., & Vingron, M. (2002). Variance stabilization applied to microarray data calibration and to the quantification of differential expression. Bioinformatics (Oxford, England), 18 Suppl 1, S96–S104. 10.1093/bioinformatics/18.suppl_1.s96

55. Ritchie, M. E., Phipson, B., Wu, D., Hu, Y., Law, C. W., Shi, W., & Smyth, G. K. (2015). limma powers differential expression analyses for RNA-sequencing and microarray studies. Nucleic acids research, 43(7), e47. 10.1093/nar/gkv007

56. Wu, T., Hu, E., Xu, S., Chen, M., Guo, P., Dai, Z., Feng, T., Zhou, L., Tang, W., Zhan, L., Fu, X., Liu, S., Bo, X., & Yu, G. (2021). clusterProfiler 4.0: A universal enrichment tool for interpreting omics data. Innovation (Cambridge (Mass.)), 2(3), 100141. 10.1016/j.xinn.2021.100141

57. Kanehisa, M., Furumichi, M., Sato, Y., Kawashima, M., & Ishiguro-Watanabe, M. (2023). KEGG for taxonomy-based analysis of pathways and genomes. Nucleic acids research, 51(D1), D587–D592. 10.1093/nar/gkac963

58. Liberzon, A., Birger, C., Thorvaldsdóttir, H., Ghandi, M., Mesirov, J. P., & Tamayo, P. (2015). The Molecular Signatures Database (MSigDB) hallmark gene set collection. Cell systems, 1(6), 417–425. 10.1016/j.cels.2015.12.004

59. Luo, W., & Brouwer, C. (2013). Pathview: an R/Bioconductor package for pathway-based data integration and visualization. Bioinformatics (Oxford, England), 29(14), 1830–1831. 10.1093/bioinformatics/btt285

60. Gu, Z., Eils, R., & Schlesner, M. (2016). Complex heatmaps reveal patterns and correlations in multidimensional genomic data. Bioinformatics (Oxford, England), 32(18), 2847–2849. 10.1093/bioinformatics/btw313

