## Supplemental data for "Polymer of methyl malonic acid suppress inflammation by downregulating IL-2 through ROS overproduction"

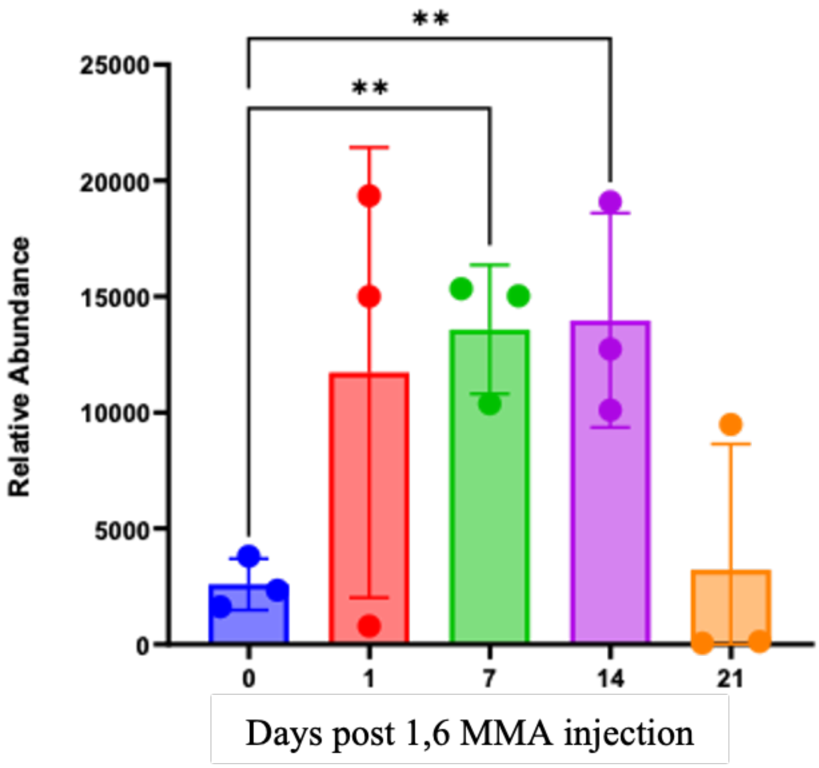


**Supplementary figure 1: Sustained release of MMA in blood post polymer injection.** Relative abundance correlating MMA amounts were calculated from mice blood across different time points. One-way Anova was used to calculate statistical differences where *, **, *** and **** are for p values less than 0.05, 0.01, 0.001 and 0.0001 respectively.

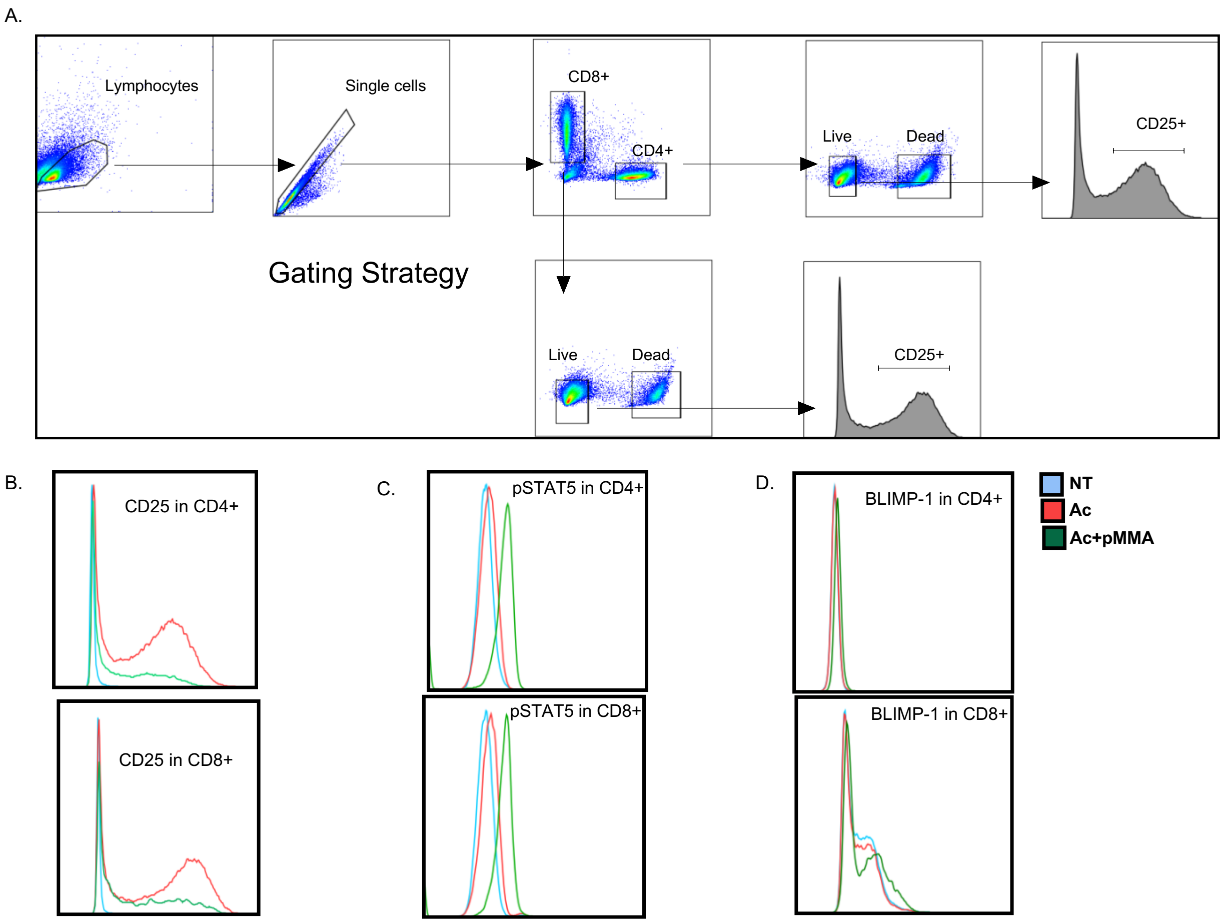


**Supplementary figure 2: Gating strategy utilized for cell surface and intracellular protein quantification using flow cytometry.** A. Flow plots were constructed to show gating strategy of CD3+ T cells to calculate frequency of live CD4+ and CD8+ t cells. Furthermore, CD25+ frequencies were calculated from these two T cell subsets. Histograms were constructed to demonstrate the difference in expression of B. CD25, C. pSTAT5 and D. BLIMP-1 in CD4+ and CD8+ between un-activated (NT) control, activated control (Ac) and activated T cells treated with 250µg/ml of 1,6 MMA.

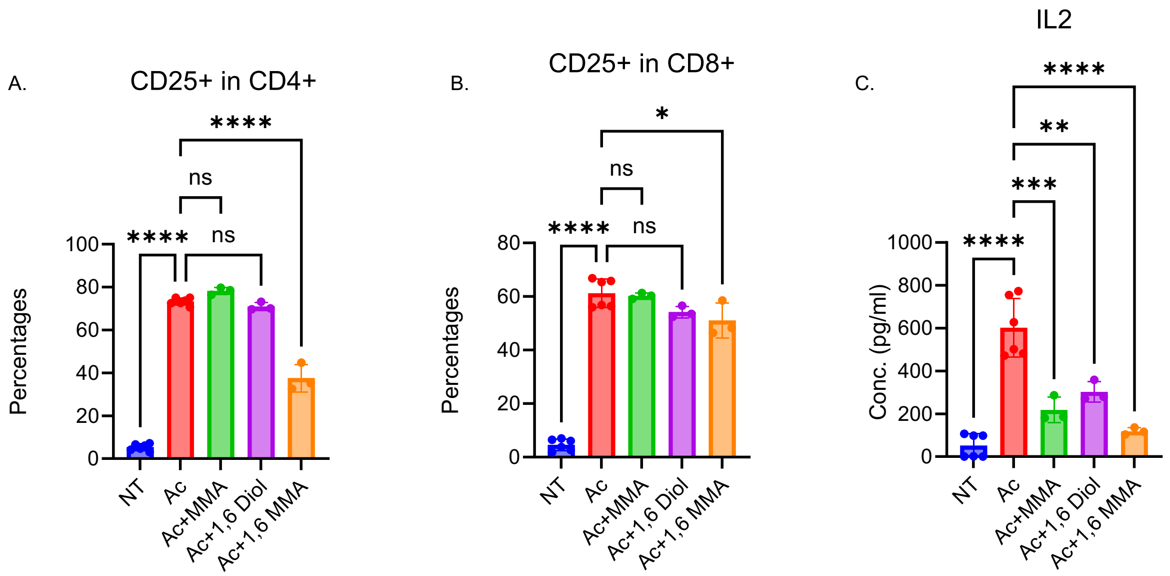


**Supplementary figure 3: Only polymer of MMA causes overall reduction in T cell activation.** CD25+ frequencies were calculated for both A. CD4+ and B. CD8+ T cells across the mentioned conditions. C. IL2 was quantified using sandwich ELISA from the cell supernatant. One-way Anova was used to calculate statistical differences where *, **, *** and **** are for p values less than 0.05, 0.01, 0.001 and 0.0001 respectively.

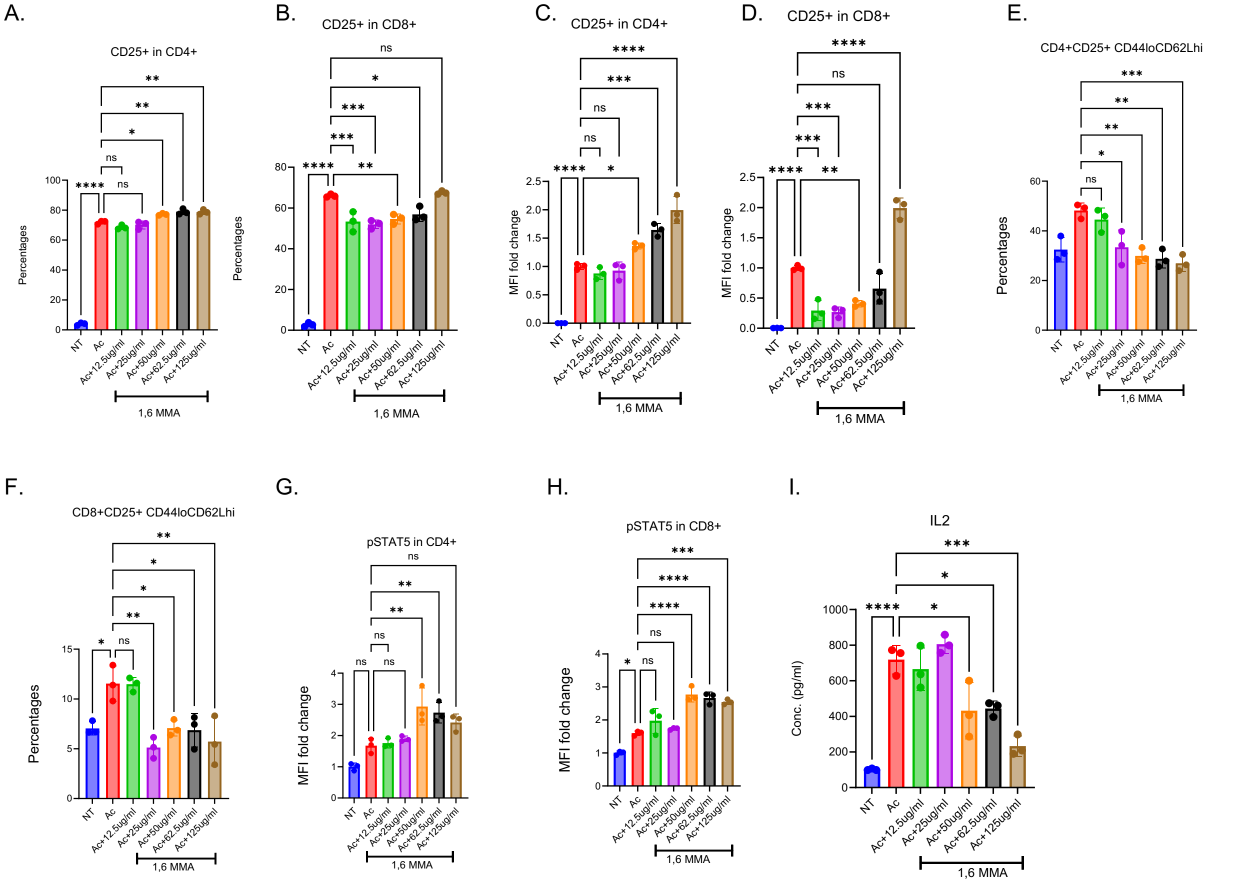


**Supplementary figure 4: Polymer of MMA reduces T cell activation in a dose dependent manner.** CD25+ frequencies (A and B) and MFI fold changes (C and D) were calculated for both CD4+ and CD8+ T cells treated with different doses of 1,6 MMA. Frequencies of E. CD4+CD25+CD44loCD62Lhi and F. CD8+CD25+CD44loCD62Lhi were calculated. pSTAT5 MFI fold changes were quantified for G. CD4+ and H. CD8+ T cells. I. Cell supernatant IL2 was quantified for the same conditions. One-way Anova was used to calculate statistical differences where *, **, *** and **** are for p values less than 0.05, 0.01, 0.001 and 0.0001 respectively.

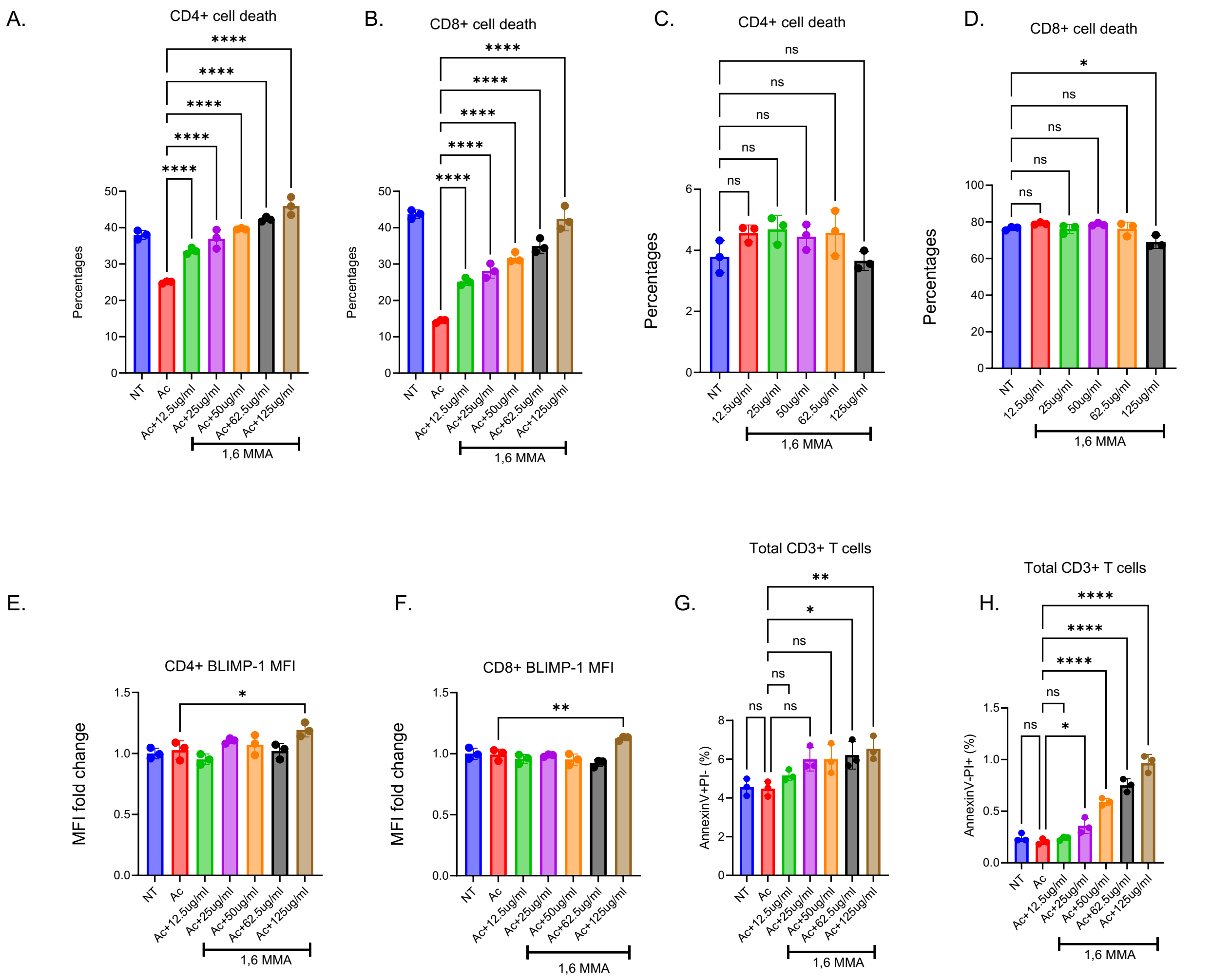


**Supplementary figure 5: Polymer of MMA enhances apoptosis in activated T cells in a dose dependent manner.** CD4+ and CD8+ T cell death was quantified using a viability dye for activated (A and B) and un-activated (C and D) T cells treated with different doses of 1,6 MMA. BLIMP-1 MFI fold changes were quantified for E. CD4+ and F. CD8+ T cells. Cells treated with different doses of 1,6 MMA were used to calculate G. AnnexinV+PI- (early apoptosis) and H. AnnexinV-PI+ (necrosis). One-way Anova was used to calculate statistical differences where *, **, *** and **** are for p values less than 0.05, 0.01, 0.001 and 0.0001 respectively.

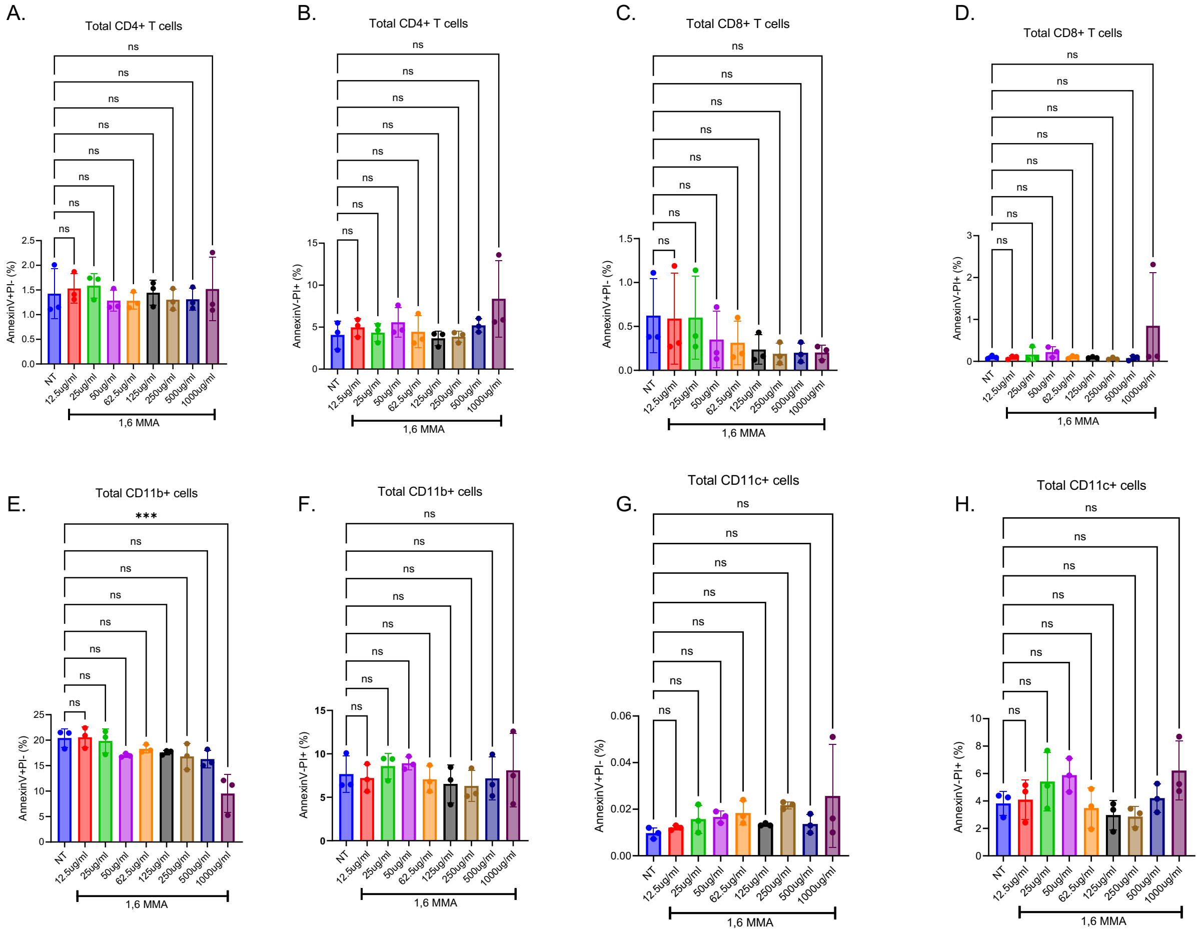


**Supplementary figure 6: Polymer of MMA does not cause non-specific cell death.** AnnexinV+PI- (early apoptosis) and AnnexinV-PI+ (necrosis) percentages were calculated for CD4+ (A and B), CD8+ (C and D), CD11b+ (E and F) and CD11c+ (G and H) for splenocytes treated with different doses of 1,6 MMA in the absence of any activating stimulus. Cells were then gated for the different immune cell populations. One-way Anova was used to calculate statistical differences where *, **, *** and **** are for p values less than 0.05, 0.01, 0.001 and 0.0001 respectively.

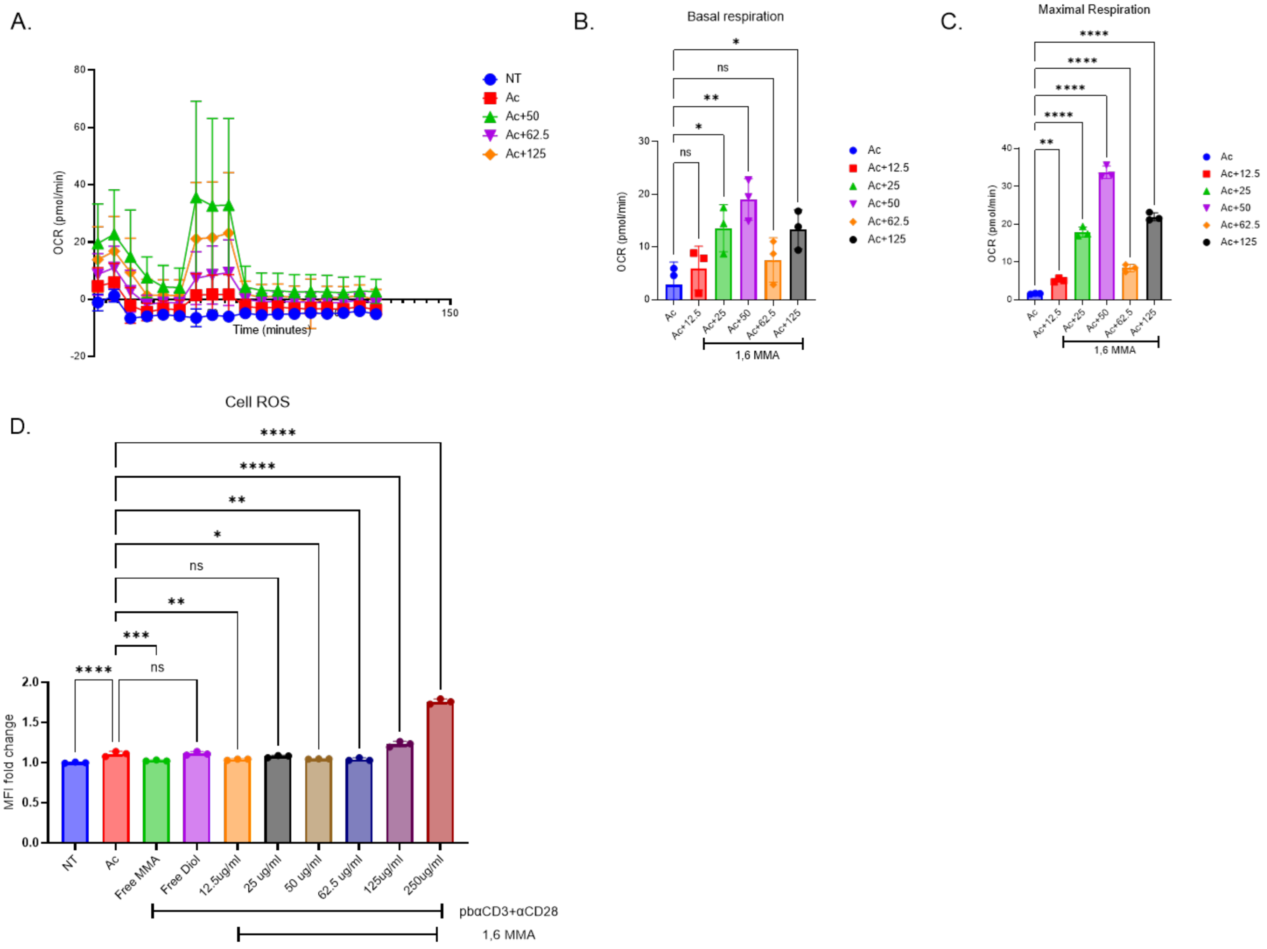


**Supplementary figure 7: T cells treated with 1,6 MMA show higher metabolic respiration with increase in ROS.** CD3+ T cells were activated in-vitro and treated with different doses of 1,6 MMA were used for A. seahorse metabolic profile analysis (OCR) from which B. basal and C. maximal respiration were calculated across the mentioned conditions. D. Flow cytometry was then used to quantify the intracellular ROS amounts. One-way Anova was used to calculate statistical differences where *, **, *** and **** are for p values less than 0.05, 0.01, 0.001 and 0.0001 respectively.

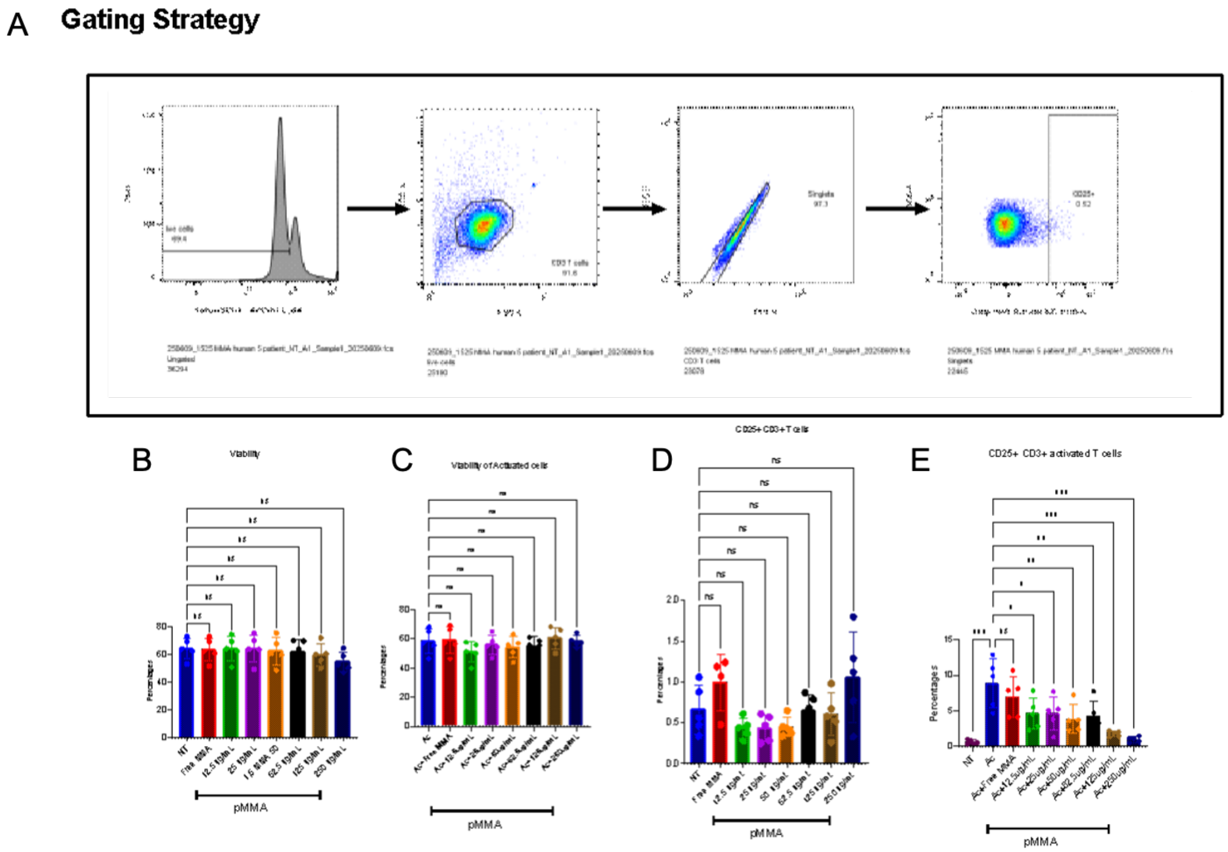


**Supplementary Figure 8. Flow cytometric gating strategy and quantification of T-cell activation across 1,6 MMA (denoted as pMMA) treatment conditions.** A. Gating strategy, cellular viability (left), followed by lymphocyte forward- and side-scatter characteristics, doublet exclusion (FSC-A vs FSC-H), and final selection of $CD3^+$ T cells. Gates are indicated by dashed boxes and arrows illustrate the gating progression. This strategy was applied uniformly across all samples and conditions. B. Quantification of cell viability across untreated (NT), MMA and pMMA-treated conditions. No statistically significant differences were observed across conditions, indicating that pMMA treatment does not impact overall cell viability. C. Viability of activated T cells, defined as the fraction of viable cells within the activated $CD3^+$ gate. Consistent viability across conditions confirms that observed differences in activation markers are not driven by cell death. D. Percentage of $CD25^+$ cells within the total non-activated $CD3^+$ T-cell population. E. Percentage of $CD25^+$ activated $CD3^+$ T cells in activated T cells show a dose-dependent reduction in CD25 expression with increasing pMMA concentration. For all bar plots, individual dots represent biological replicates, bars denote mean ± SEM, and statistical comparisons were performed as described in Methods. Significance is indicated as $p < 0.05$ (*), $p < 0.01$ (**), $p < 0.001$ (***), or not significant (ns).

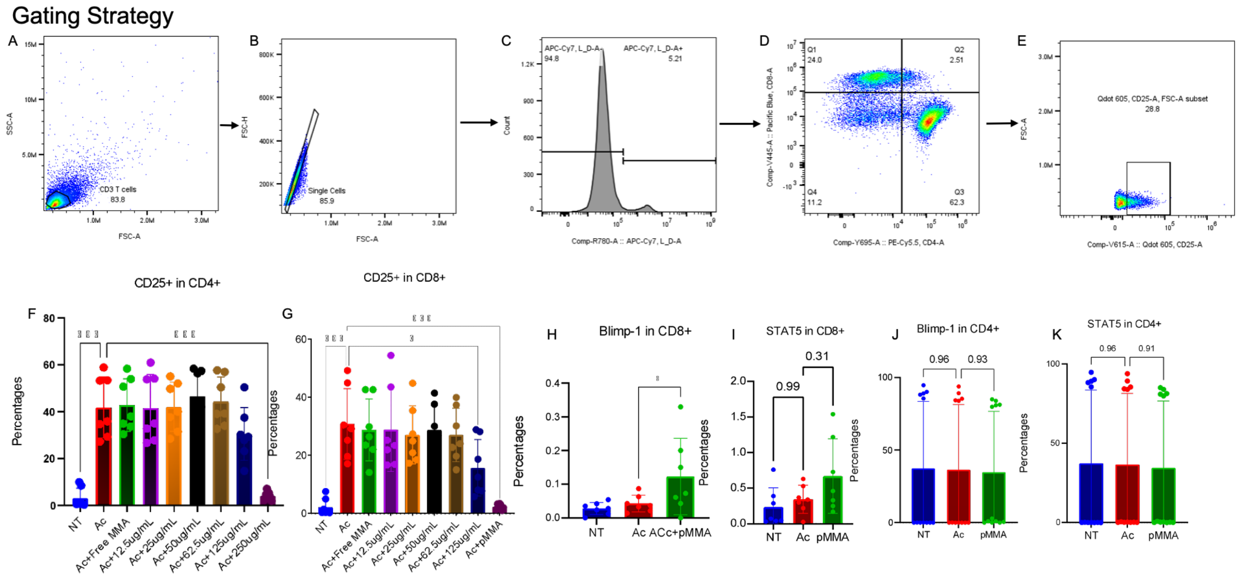


**Supplementary Figure 9. Flow cytometric gating strategy and CD25, BLIMP-1 and STAT5 expression in CD4⁺ and CD8⁺ T cells across 1,6 MMA (shown as pMMA) conditions.** (A) - (E) Flow cytometry gating strategy. Events were sequentially gated on forward- and side-scatter properties to exclude debris (A), followed by doublet exclusion using FSC-A versus FSC-H B. Live cells were identified using a viability dye (C), after which $CD3^+$ T cells were gated and subdivided into $CD4^+$ and $CD8^+$ populations (D). Within these populations, $CD25^+$ cells were identified using FSC-A–restricted gates (E). Percentages within each gate are indicated. The same gating strategy was applied uniformly across all samples. F. Percentage of $CD25^+$ cells within the $CD4^+$ T-cell across untreated (NT), activated (Ac), MMA and increasing pMMA concentrations. G. Percentage of $CD25^+$ cells within the $CD8^+$ T-cell under the same conditions. (H) Intracellular expression of Blimp-1 in $CD8^+$ T cells. I. Intracellular STAT5 levels in $CD8^+$ T cells. J. Intracellular Blimp-1 expression in $CD4^+$ T cells. K. Intracellular STAT5 expression in $CD4^+$ T cells. For all quantitative panels, individual dots represent biological replicates, bars indicate mean ± SEM, and statistical comparisons were performed as described in Methods. Significance values are shown above comparisons; non-significant differences are explicitly indicated.

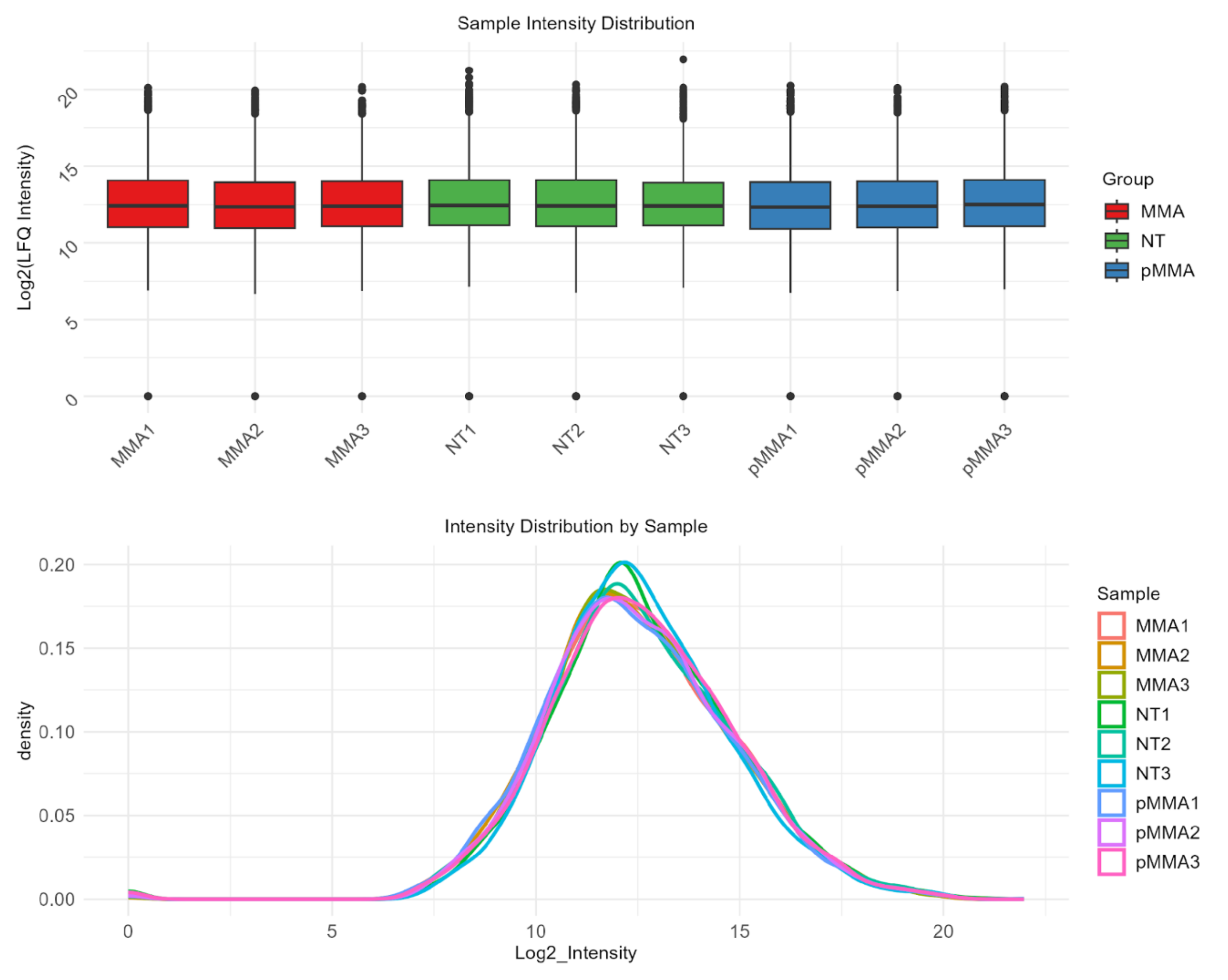


**Supplementary Figure 10. Quality control assessment of LFQ proteomics data across experimental conditions.** Boxplot of $\log_2$-transformed LFQ intensity distributions for individual samples across NT, MMA, and 1,6 MMA (pMMA) conditions. Each box represents the interquartile range (IQR) with the median indicated by the central line; whiskers denote 1.5× IQR, and points represent outliers. Comparable median intensities and distribution widths across samples indicate consistent overall signal intensity and absence of global batch effects prior to downstream normalization and differential expression analysis. (b) Density plots of $\log_2$ LFQ intensity distributions for each individual sample. Overlapping density curves demonstrate highly similar global intensity profiles across biological replicates and experimental conditions, supporting uniform proteome coverage and technical reproducibility. Minor variation at the distribution tails reflects expected biological and measurement variability rather than systematic bias.

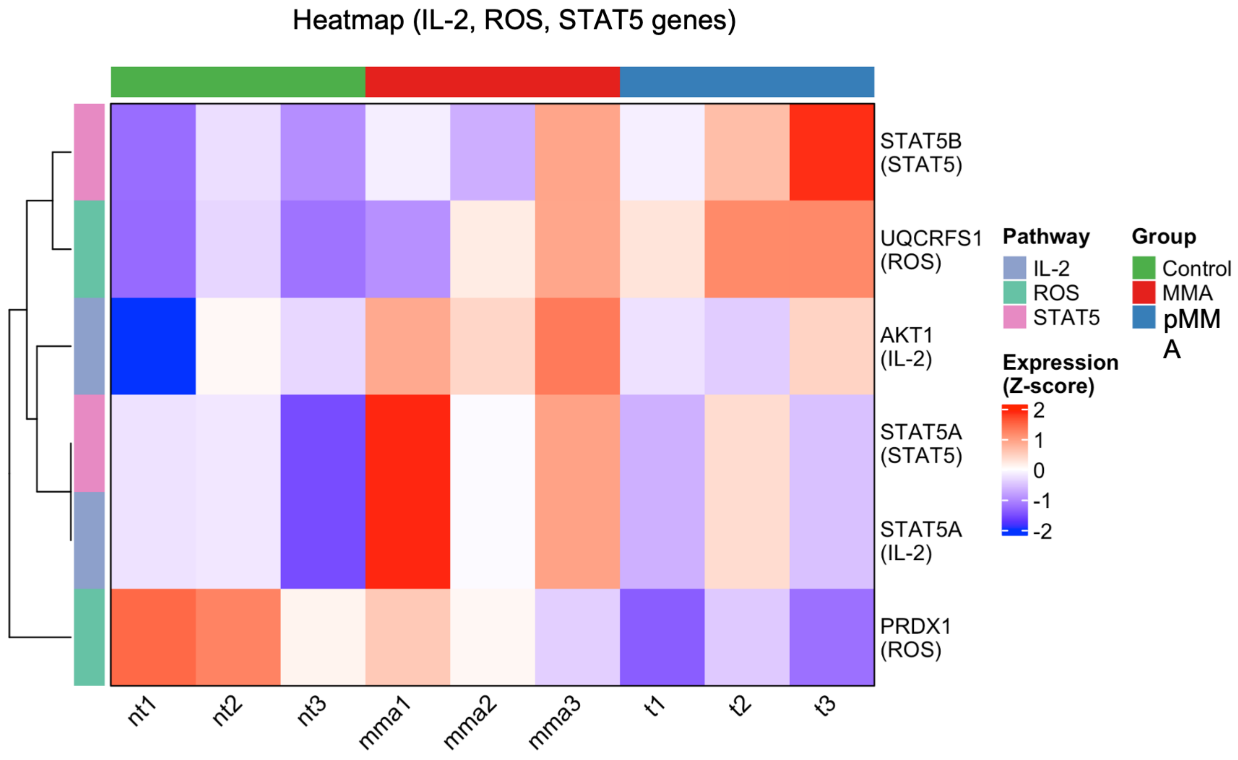


**Supplementary Figure 11. Differential expression of IL-2, ROS, and STAT5 pathway–associated genes across experimental conditions.** Heatmap depicting z-score–normalized $log_2$ LFQ expression levels of selected proteins associated with IL-2 signaling, reactive oxygen species (ROS) regulation, and STAT5 signaling across NT (control), MMA, and 1,6 MMA (pMMA) conditions. Each column represents an individual biological replicate, and each row corresponds to a protein, annotated by its primary functional pathway. Hierarchical clustering was performed on rows using Euclidean distance, grouping proteins with similar expression patterns across conditions. The color scale indicates relative expression changes (blue, lower than mean; red, higher than mean), enabling comparison across samples independent of absolute abundance.
